# Somatic chromosomal number alterations affecting driver genes inform *in-vitro* and clinical drug response in high-grade serous ovarian cancer

**DOI:** 10.1101/2020.10.04.325365

**Authors:** Filipe Correia Martins, Dominique-Laurent Couturier, Ines de Santiago, Carolin Margarethe Sauer, Maria Vias, Mihaela Angelova, Deborah Sanders, Anna Piskorz, James Hall, Karen Hosking, Anumithra Amirthanayagam, Sabina Cosulich, Larissa Carnevalli, Barry Davies, Tom B. K. Watkins, Gabriel Funingana, Helen Bolton, Krishnayan Haldar, John Latimer, Peter Baldwin, Robin Crawford, Matthew Eldridge, Bristi Basu, Mercedes Jimenez-Linan, Nicholas McGranahan, Kevin Litchfield, Sohrab P. Shah, Iain McNeish, Carlos Caldas, Gerard Evan, Charles Swanton, James D. Brenton

## Abstract

The genomic complexity and heterogeneity of high-grade serous ovarian cancer (HGSOC) has hampered the realisation of successful therapies and effective personalised treatment is an unmet clinical need. Here we show that primary HGSOC spheroid models can be used to predict drug response and use them to demonstrate that somatic copy number alterations (SCNAs) in frequently amplified HGSOC cancer genes significantly correlate with gene expression and drug response. These genes are often located in areas of the genome with frequent clonal SCNAs. MYC chromosomal copy number is associated with ex-vivo and clinical response to paclitaxel and ex-vivo response to mTORC1/2 inhibition. Activation of the mTOR survival pathway in the context to MYC-amplified HGSOC is mostly due to increased prevalence of SCNAs in genes from the PI3K pathway. These results suggest that SCNAs encompassing driver genes could be used to inform therapeutic response in the context of clinical trials testing personalised medicines.

## Main

Progressing precision medicine for high grade serous ovarian carcinoma (HGSOC) has been significantly impeded by its highly heterogeneous nature, driven by chromosomal instability (CIN) resulting in divergent evolution (1, 2). Homologous recombination deficiency (HRD) is the commonest actionable mutational state, notably from germline or somatic deleterious *BRCA1* and *BRCA2* mutations, and routine use of PARP maintenance therapy has significantly extended progression-free survival(3, 4). PARP therapy targets a loss of function phenotype exploiting synthetic lethality but biomarker-driven approaches to identify and target gain of function drivers have not been clinically developed.

The frequency of nucleotide substitutions that are “actionable” in HGSOC is ∼1% and the vast majority of genomic changes are structural variants, with high frequency of somatic chromosomal number alterations (SCNAs). Recurrent patterns of SCNAs can be identified across multiple tumours(5, 6) as a result of distinct ordering throughout tumour evolution or parallel selection (6-8). Multiregional analysis across 22 tumour types revealed frequent subclonal focal amplifications in chromosomes 1q (encompassing *BCL9* and *MCL1*), 5p (*TERT*), 11q (*CCND1*), 19q (*CCNE1*) and 8q (*MYC*). *MYC, CCNE1, TERT, KRAS* and genes from the PI3K/AKT/mTOR pathway (eg. *PIK3CA*) are amongst the commonest amplified cancer genes in HGSOC (2, 9-13).

SCNAs frequently drive gene expression and protein levels (14, 15) and detection of ERBB2 amplification in breast carcinoma is a critically important biomarker for trastuzumab therapy (16, 17). Bulk gene expression profiling has not provided strong evidence of driver gene expression owing to common confounding effects from other cells in the tumour microenvironment (18-20). By contrast, use of bulk sequencing to detect SCNA, even in low to moderate cellularity cancer specimens, retains high specificity. In HGSOC, SCNA can be efficiently detected using shallow whole genome sequencing from clinical biopsies (21-24).

We hypothesised that SCNA characterisation in HGSOC might provide specific and clinically relevant biomarkers for molecularly targeted therapy. Amongst the different cell culture and animal models developed to interrogate HGSOC biology (25-29), studies using a limited number of HGSOC organoids and spheroids (cultured and uncultured clusters of primary ovarian cancer cells from HGSOC patient ascites, respectively) suggested that they could be used to predict platinum resistance (29, 30). We here show how spheroids represent the global HGSOC genomic diversity and use them to assess how *in-vitro* drug response correlates with clinical response and SCNAs affecting putative HGSOC driver genes (Figure 1, Supplementary Figure 1 and Table 1).

**Table 1.**
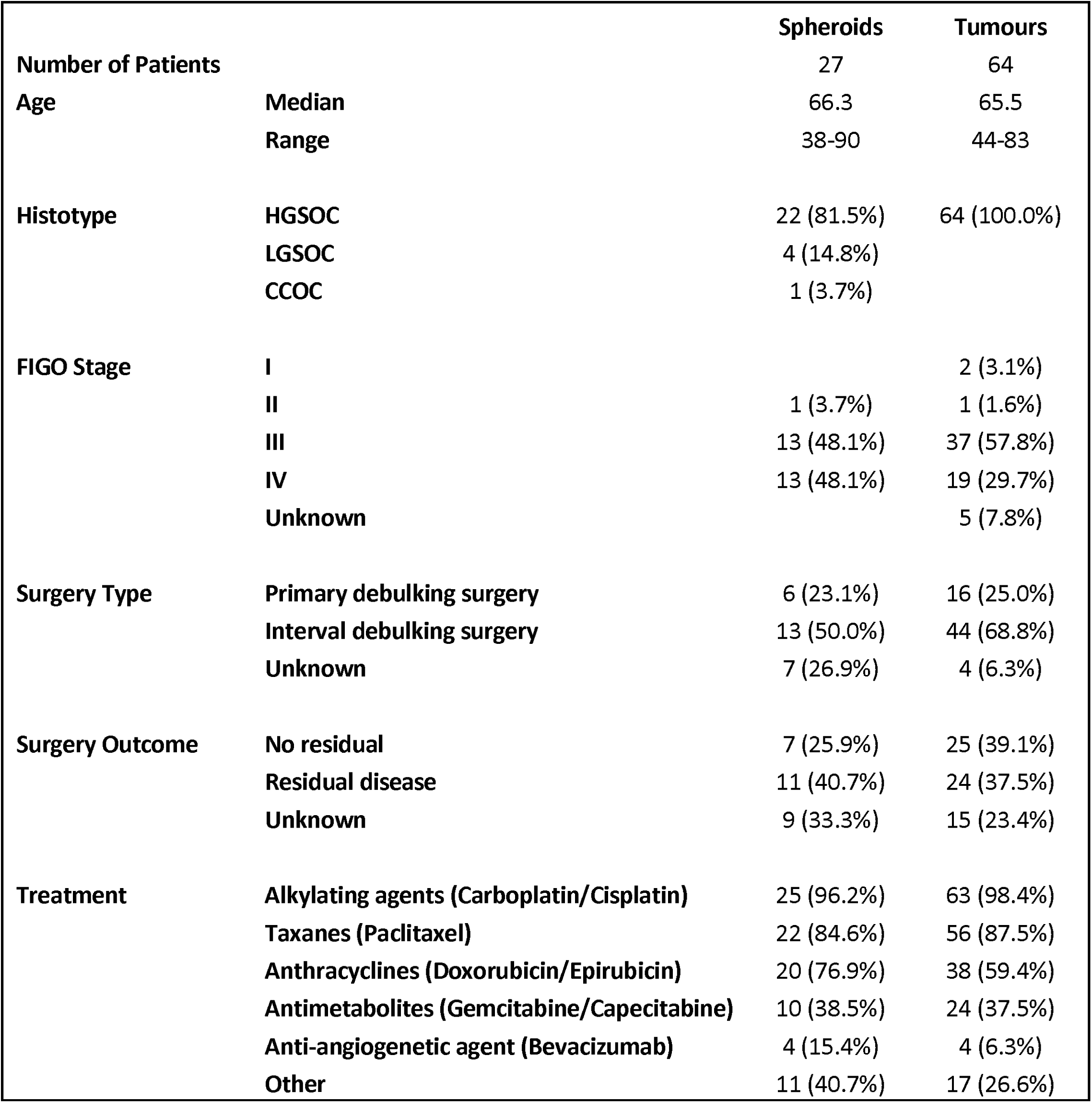
Demographic characterisation of the OVO4 clinical cohort

**Figure 1.**
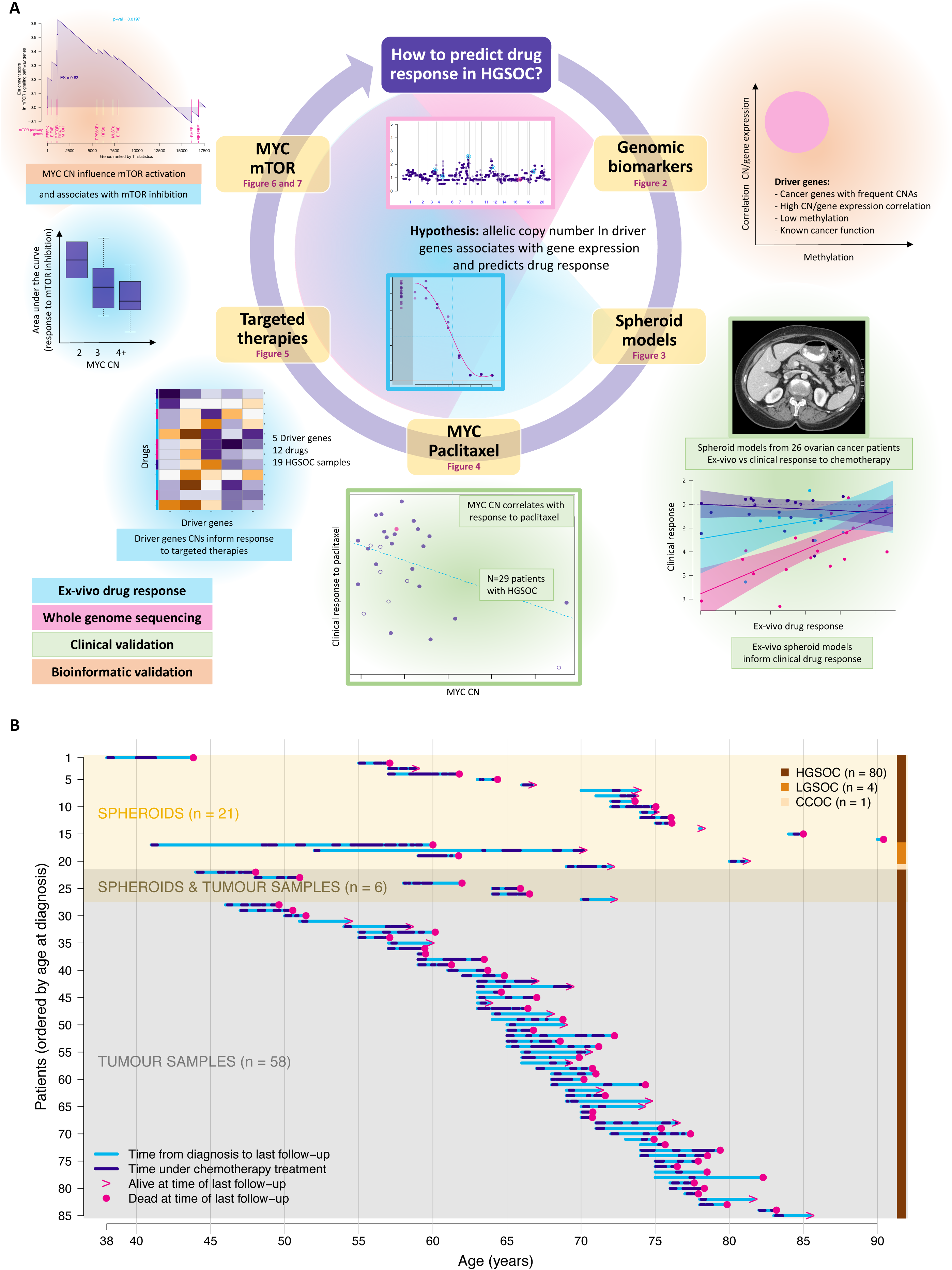
A. Summary of study. graphically representing question, aims and methods used. **B.** Timeline with characterisation of the OVO4 clinical cohort used in this study and time of follow-up for each patient with information related to patient status at last follow-up, chemotherapy treatment period(s) and cancer type.

Recent clinical trials using combinations of chemotherapeutic agents, PARP inhibitors and/or other targeted therapies in HGSOC and other CIN-driven tumours showed effective response in a subset of patients(31-33). Mutational characterisation of the tumours did not distinguish responders from non-responders. Our work demonstrate that SCNAs in putative driver genes inform how each HGSOC responds to individual targeted therapies and that sWGS is a crucial tool to predict response in the context of clinical trials for CIN-driven tumours.

## Results

### HGSOC putative driver genes showed strong associations between chromosomal copy number and gene expression

In order to select a small number of copy number driver genes to test as predictors of drug response, we first examined the genomic data from the TCGA HGSOC cohort (Figure 2A; grey shadowing). The GISTIC (“Genomic Identification of Significant Targets in Cancer)” algorithm was previously used to define recurrent somatic copy number alterations (SCNAs) across cancers (34, 35). Although strong associations between chromosomal copy number and gene expression have been shown previously (14, 15), it is unclear if these associations are stronger in genes defined as cancer drivers. In the TCGA cohort, 2415 genes were affected by amplifications or homozygous deletions in more than 5% of the cases. 156/2415 (6.5%) were putative drivers based on external curation in the OncoKB Cancer Gene List (13). These putative SCNA drivers showed the highest correlation between chromosomal copy number and gene expression, when compared to non-cancer genes and cancer genes from OncoKB with <5% SCNAs (Figure 2B). They were also more frequently hypomethylated when compared with the remaining genes in TCGA HGSOC cohort (Figure 2C) and methylation was a major determinant of the correlation between chromosomal copy number and gene expression (Figure 2D).

**Figure 2.**
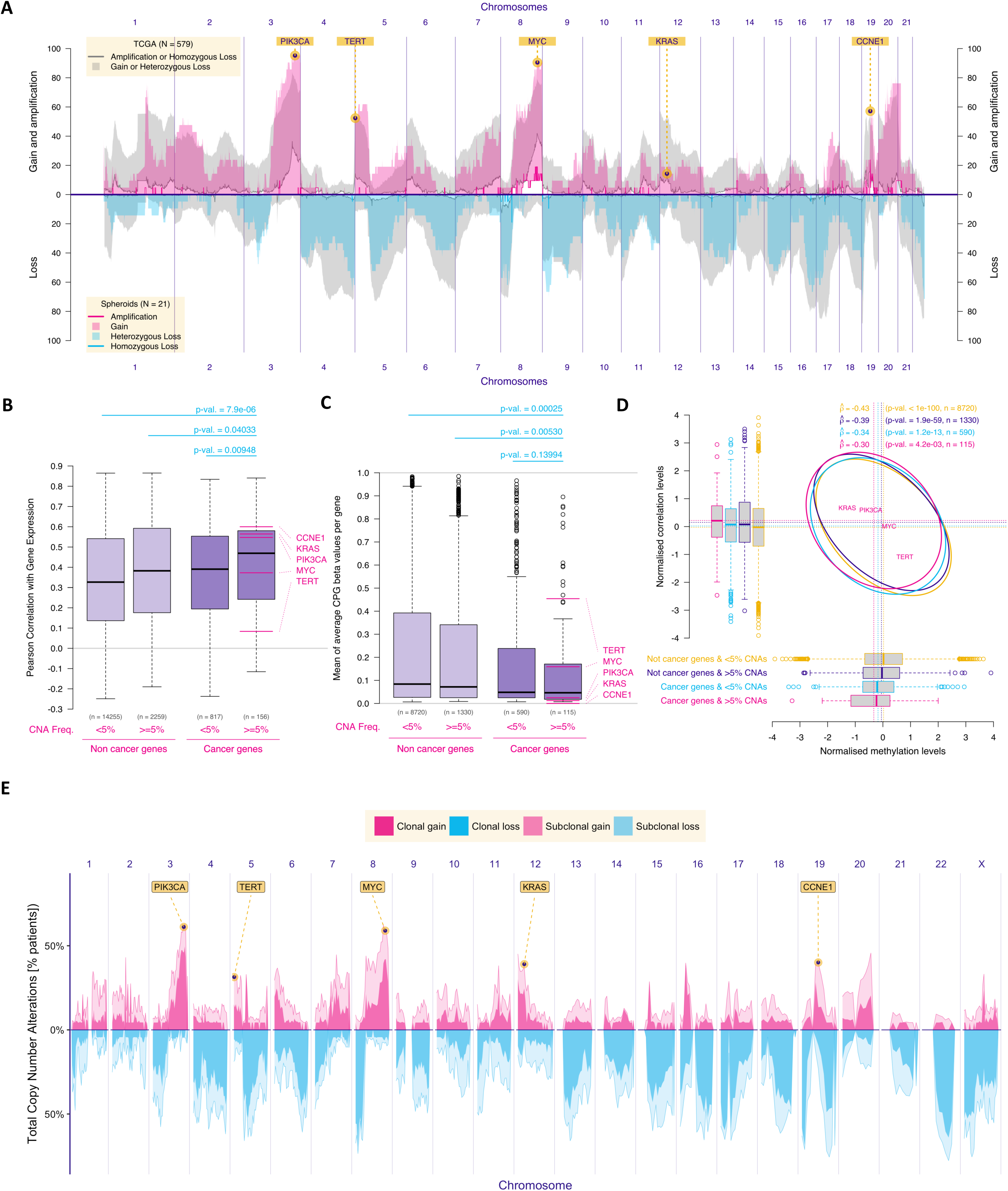
HGSOC driver genes from the TCGA cohort showed strong associations between chromosomal copy number and gene expression and were frequently affected by SCNAs in the spheroid cohort. **A**. Plot showing the prevalence of chromosomal alterations across the genome in both the TCGA cohort (n=579) and in the cohort of patients with spheroid samples (n=21). For the spheroid cohort, gain was defined as 3 or 4 chromosomal adjusted copies and amplifications as 5 or more adjusted copies. For the TCGA cohort, categories as available in the TCGA database were used **B.** Boxplots showing the Pearson’s correlation scores between gene expression and respective chromosomal copy number for each gene, split between cancer vs non-cancer genes and prevalent (>5% SCNAs in HGSOC) vs non-prevalent genes. Driver genes (in the far right; defined as ‘cancer genes’ that have SCNA alteration frequency in at least 5% of the samples) had the highest positive correlation scores. Numbers on top of the boxplots correspond to the p-values obtained with Mann– Whitney-Wilcoxon tests. **C.** Boxplots showing methylation levels (beta-values) for all genes, split in the same groups as in panel A. Prevalent non-cancer genes were significantly more methylated than prevalent cancer genes (Mann–Whitney-Wilcoxon’s p-value: 0.005). **D.** Plot showing, on the normalised scale, that methylation levels and correlation of chromosomal copy number and gene expression were associated. **E.** Frequency of somatic clonal and subclonal copy number alterations across the genome of 72 regions from 28 HGSOC primary tumours. Gains and losses were classified relative to ploidy. The plot showing the genomic distribution of the frequency of somatic copy number alterations across 127 regions of both primary tumours and metastases from 30 HGSOC patients is presented in Supplementary figure 2.

We further analysed 127 anatomically distinct HGSOC samples from a cohort of 30 cases with multi-regional sampling(36). We characterized SCNA as clonal if they were present in all regions from the same individual (median of 4 samples), and subclonal if present in at least one but not all regions. Subclonal SCNAs were extremely common and distributed across the genome (Figure 2E and Supplementary Figure 2). Focal clonal changes were located in areas of the genome with frequent SCNAs (Figures 2A and E) and involved the HGSOC drivers *PIK3CA, TERT, MYC, KRAS* and *CCNE1*(9).

### Ex-vivo drug response using human spheroids predicted clinical response to the first and second line chemotherapeutic regimens

We performed shallow WGS (sWGS) on 26 primary spheroids from women with ovarian cancer and adequate numbers of vials for functional analysis (Methods; Supplementary Figures 3A-Z). Twenty-one out of the 26 were HGSOC. In order to assess whether these spheroid samples were representative of the genomic landscape of HGSOC, we compared their frequency of SCNAs with the TCGA cohort. The distribution of spheroid SCNA was similar to the distribution of amplifications and losses from TCGA (Figure 2A).

**Figure 3.**
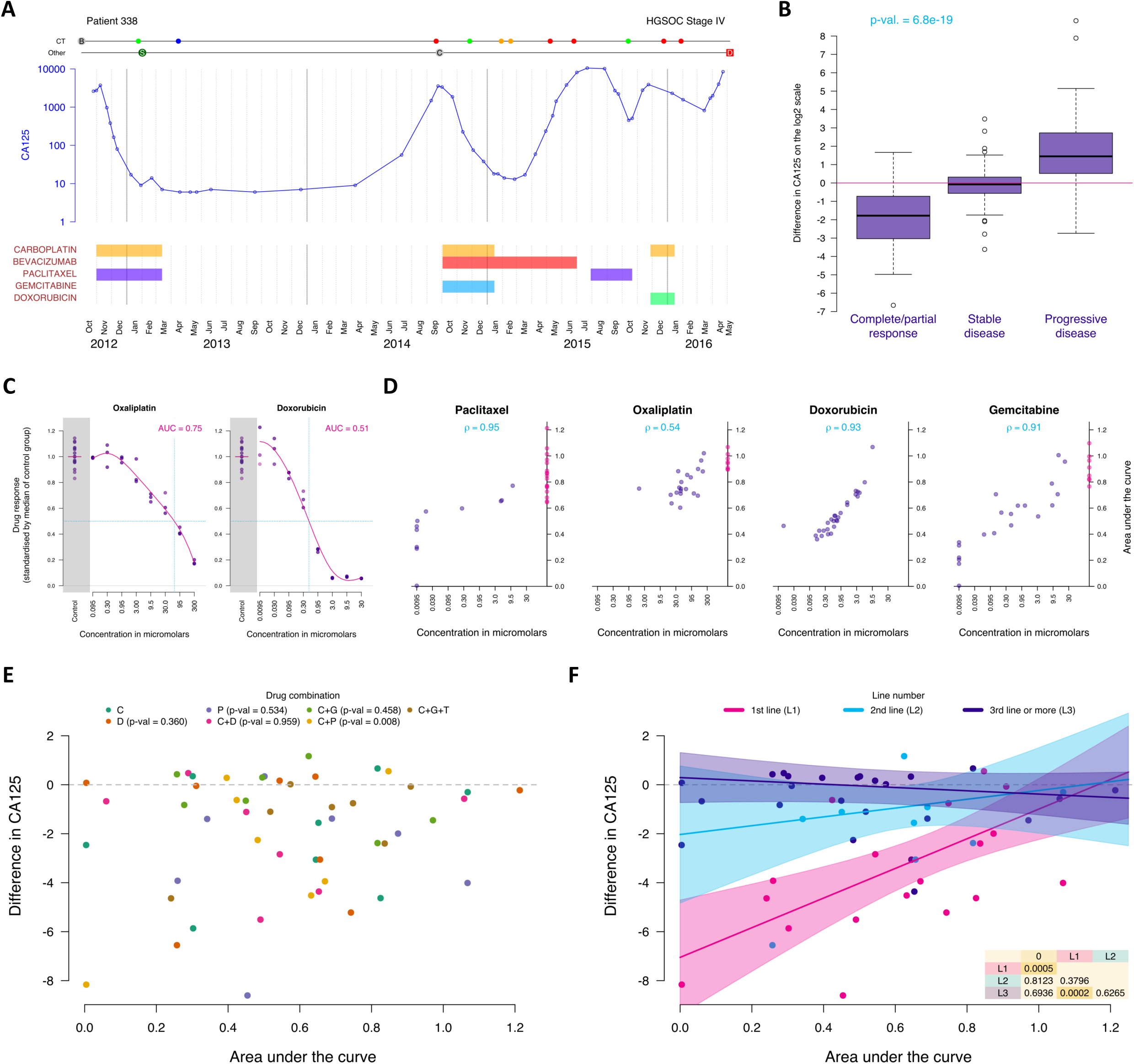
Ex-vivo drug response in primary spheroid samples is associated with clinical response to chemotherapy. **A**. Example of a patient timeline summarising the available clinical data (CA125 levels, chemotherapy cycles, CT scan results); S: surgery, C: collection, D: death; CT is represented as blue (complete response), green (partial response), yellow (stable disease) and red (progressive disease). **B.** Boxplots showing the association between variation in CA125 (measured by the difference of CA125 values in log 2 scale after and before each chemotherapy cycle) split by categories of CT radiological response and p-value of the Jonckheere-Terpstra test of association. **C.** Example of *in vitro* response to oxaliplatin and doxorubicin in a spheroid sample. Each dot measures cell viability against the average cell viability of the controls for each drug concentration after 5 days incubation. The pink line corresponds to the M-spline fit. AUC was obtained by integrating the drug response M-spline fit over the range of interest on the log scale. Intersection of dotted blue lines indicates IC50. Dot plot in grey shading indicates control data. **D.** Scatterplots showing the association between area under the curve values and IC50 for selected drugs. Each dot represents one sample. Pink dots represent cases where IC50 was not reached owing to viability ≤50% at maximum dose Spearman’s Rho correlation estimates are indicated in turquoise. **E & F**. Scatterplots showing the clinical responses measured by variation in logarithmic scale of CA125 during each chemotherapy regimen/line in each patient (y axis) as a function of the *in-vitro* response to the same drugs, measured by area under the curve on samples from the same patient (x axis). When combinations of drugs were used in the clinical setting, the combined AUC was obtained by multiplying AUC for individual drugs; Points are colour-coded by drugs (panel E) or chemotherapy line numbers (panel F). In panel E, the p-values of the Wald t-tests defining if the slope parameter of linear mixed models corresponding to each drug are different from 0 are indicated in the legend. In panel F, the mixed model fitted lines per chemotherapy lines and corresponding 95% confidence intervals (shaded areas) are indicated. The p-values of the Wald t-tests performing pairwise comparison of the chemotherapy line slopes or comparing the chemotherapy line slopes to 0 are indicated P: paclitaxel; C: carboplatin; D: doxorubicin; G: gemcitabine; T: targeted inhibitor.

To test the performance of *in vitro* drug response from spheroids, we compared it with the clinical response in patients from whom spheroids were derived. Clinical response measures have been previously categorized semi-quantitatively (30, 37). In order to define a continuous variable that integrated all parameters of the degree of response, we compared variation in serum Cancer Antigen 125 (CA125) levels with radiological response on CT scans during treatment, as surrogates of histological response to chemotherapy (Supplementary Figures 3A-Z) (38). The difference between levels of CA125 levels measured in log2 scale at the time of CT imaging had the best performance as a numerical predictor of CT-inferred variation in the disease burden (P ≪0.001; Figures 3A and 3B and Supplementary Figure 4). We quantified cell viability in 26 primary human-derived spheroid samples after 5 days of ex vivo exposure to eight concentrations of each individual drug. The responses to four standard of care drugs (oxaliplatin, paclitaxel, doxorubicin and gemcitabine; Figure 3C) were compared to drug-free control spheroid samples: these were reproducible across technical and biological replicates and were not influenced by the length of the drug viability assay. Oxaliplatin was substituted for cisplatin and carboplatin as these are inactivated by DMSO and produces highly similar cellular effects (39). Area under the curve values (AUC; Fig.3C) were well correlated with IC50 estimates in drug response plots (Figure 3D). When we compared those estimates to the observed clinical response in the same patient to the same drug or class of drugs, we found that the correlation between *in vitro* and clinical response was strongest for the combination of platinum-based and paclitaxel therapy compared to other combinations or isolated platinum or paclitaxel (Figure 3E). Considering that this combination is the standard first-line treatment for HGSOC, we assessed if the response correlation was associated with the lines of treatment and found that it is strongest in the first and second line (Figure 3F).

**Figure 4.**
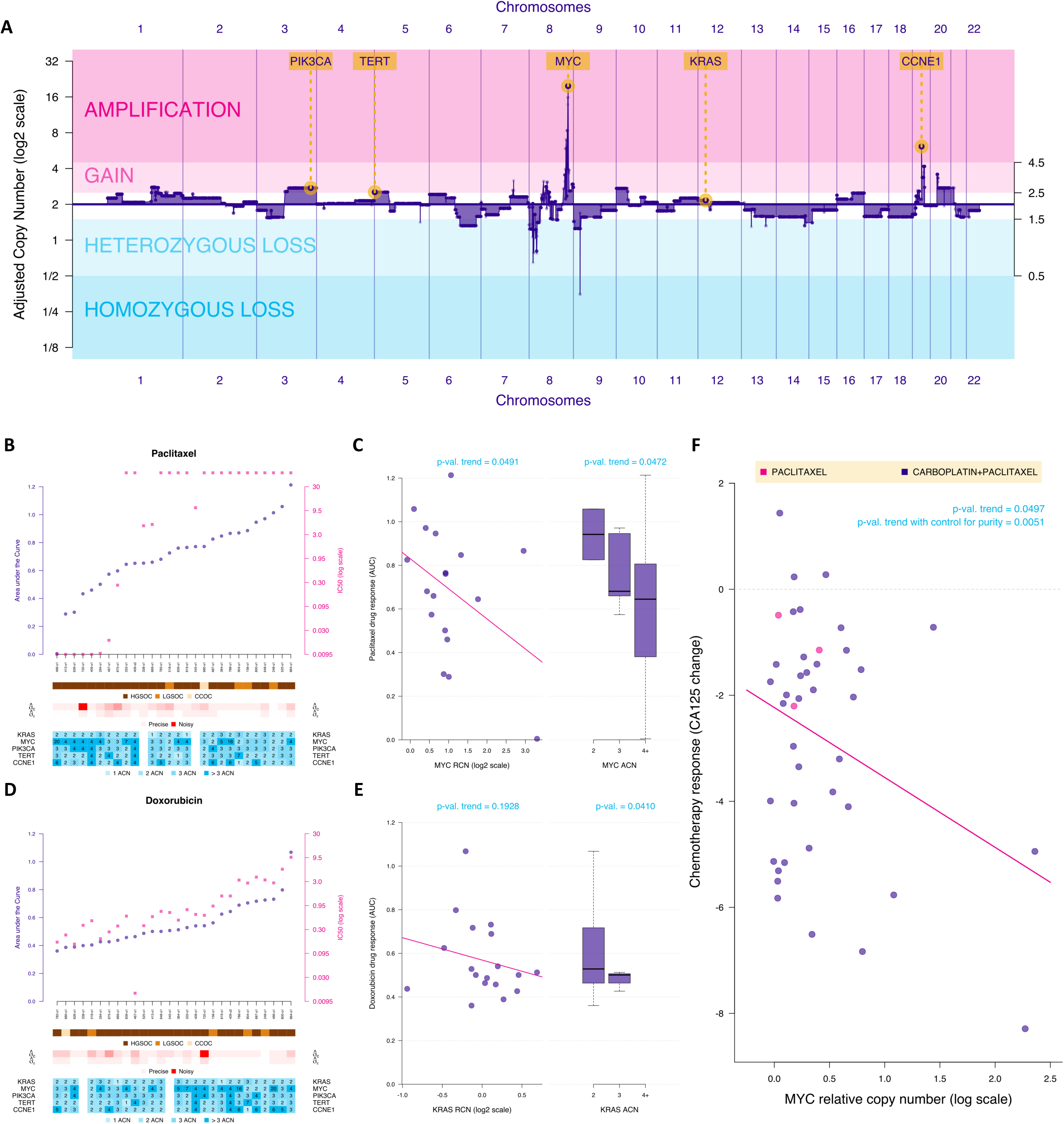
MYC amplification is a clinical biomarker of response to paclitaxel. **A**. Plot showing an example of genomic profile (adjusted copy number for each gene) for one HGSOC sample (patient 466). **B.** Scatterplot showing paclitaxel response measured by AUC (purple; left y-axis) and IC50 (pink; right y-axis) for all samples ordered by AUC levels. In the lower bars, we show, for each sample, the histological diagnosis (HGSOC – high-grade serous; LGSOC – low-grade serous; CCOC – clear cell), the normalised copy number for MYC, PIK3CA, KRAS, CCNE1 and TERT and the ex-vivo variability in the control and experimental conditions (sigma-c and sigma-e, respectively). Cases with sigma-c or sigma-e above 0.5 were considered too “noisy” and removed from the analysis in the plots 4C and E. **C.** Scatterplot and boxplots showing the associations between response to paclitaxel *in vitro* (measured by AUC) and MYC relative copy-number (RCN; left plot) and normalised absolute copy-number (3-level ACN; defined by the absolute numbers normalised for a diploid genome, to allow comparisons; right plot) P-values corresponding to the presence of a trend (linear model Wald t-test on the left and Jonkhere-test on the right) are indicated. **D.** Scatterplot showing doxorubicin response measured in all samples following the same format as in panel B. **E.** Association between KRAS RCN and ACN and doxorubicin *in vitro* response (as in panel C). **F.** Scatterplot showing the clinical response to paclitaxel (alone and in combination with carboplatin) as a function of MYC RCN. The linear mixed model p-value for the trend obtained with or without correcting for are indicated (refer to the method section for detail).

### MYC copy number is a predictor of response to paclitaxel both in the ex-vivo spheroid models and in the clinical setting

Previous genome-wide siRNA screens identified Myc as a paclitaxel sensitizer (40). Therefore, we tested the hypothesis that the number of MYC copies correlated with response to paclitaxel and found that higher number of MYC copies was associated with better response to paclitaxel using the spheroid models (p-value<0.05; Figures 4B and C). Previous work has shown that topoisomerase inhibition is synthetically lethal in the context of KRAS mutant colorectal cell lines (41). Gain of KRAS in our HGSOC samples was associated with better response to doxorubicin in spheroid models (Figure 4D and E). We then investigated whether these findings could be translated to the clinical setting and showed a similar correlation between relative copy-number of MYC and magnitude of CA125 change in response to carboplatin/paclitaxel, respectively, suggesting that MYC amplification could be a predictive clinical biomarker of response to those drugs (p-value: 0.005; Figure 4F). Considering that the number of patients receiving doxorubicin in the first or second line of chemotherapy is small and the prevalence of KRAS gain and amplification is low (Supplementary Figures 3A-Z), we were unable to validate the potential KRAS-doxorubicin association in the clinical setting.

### Copy number of putative driver genes informs response to targeted therapies

In order to further test the hypothesis that SCNAs affecting specific driver genes influence both tumour behaviour and response to molecularly targeted agents, we correlated the response to drugs targeting important regulatory genes in PI3K pathway, cell cycle and DNA repair mechanisms with the SCNAs in frequent HGSOC drivers (MYC, PIK3CA, KRAS, TERT and CCNE1; Figure 2). First, we found that, despite the significant heterogeneity of response to the different drugs used, there was a group of spheroid samples that responded similarly to all the inhibitors targeting the PI3K pathway (Figure 5A). We therefore did a pairwise comparison between responses to individual drugs. We found that response to inhibitors of the PI3K pathway correlated better between them than when compared to response to other drugs, suggesting that targeting the nodal points from the same pathway tend to have a similar effect on viability of HGSOC spheroids (p-value: 0.0001; Figure 5B). We then assessed how driver gene SCNAs correlated with response to specific targeted therapies and showed that previous established knowledge on these driver genes supported the correlations with drug response observed (Figure 5C). For example, CCNE1 amplification is frequently present in tumours with competent HR pathways and has been associated with chemotherapy resistance (42). In the context of HR deficiency, where co-existing CCNE1 amplification is uncommon (43), the ATM pathway is frequently activated. Our spheroid model not only showed that CCNE1 amplification was associated with platinum-resistance but also that lower CCNE1 copy number spheroids respond better to ATM inhibition (AZD0156; Figures 5C-E). In line with previous evidence suggesting cross-talk between mitogenic Ras/MAPK and survival PI3K/AKT pathways(44), our data also suggested that KRAS copy number positively correlates with response to AKT inhibition (AZD5363; Figure 5C). Additionally, signalling via the mTOR pathway has also been shown to regulate translational and post-translational telomerase activity(45) and our data demonstrated that TERT-amplified HGSOC samples were susceptible to inhibition of PIK3CA, AKT or mTOR (Figures 5C). Finally, inhibition of mTOR has been shown to be lethal in MYC-driven haematological tumours. In a previous phase 1 clinical trial combining paclitaxel and the dual mTORC1/2 inhibitor vistusertib (AZD2014) as a therapeutic strategy for HGSOC, there was a patient who experienced a complete response in their measurable lesions. The tumour lesions in this patient harboured a MYC mutation (33) in the Myc homology-box 2 (MBII), which is a highly preserved region across species and myc isoforms (46, 47). We therefore assessed the effect of dual m-TORC1/2 inhibition in HGSOC spheroids with MYC-amplification and showed that they were more sensitive to this targeted inhibition than tumour samples with neutral copy number (Figures 5F and G, Welch p-value:0.02; Jonkhere p-value:0.08).

**Figure 5.**
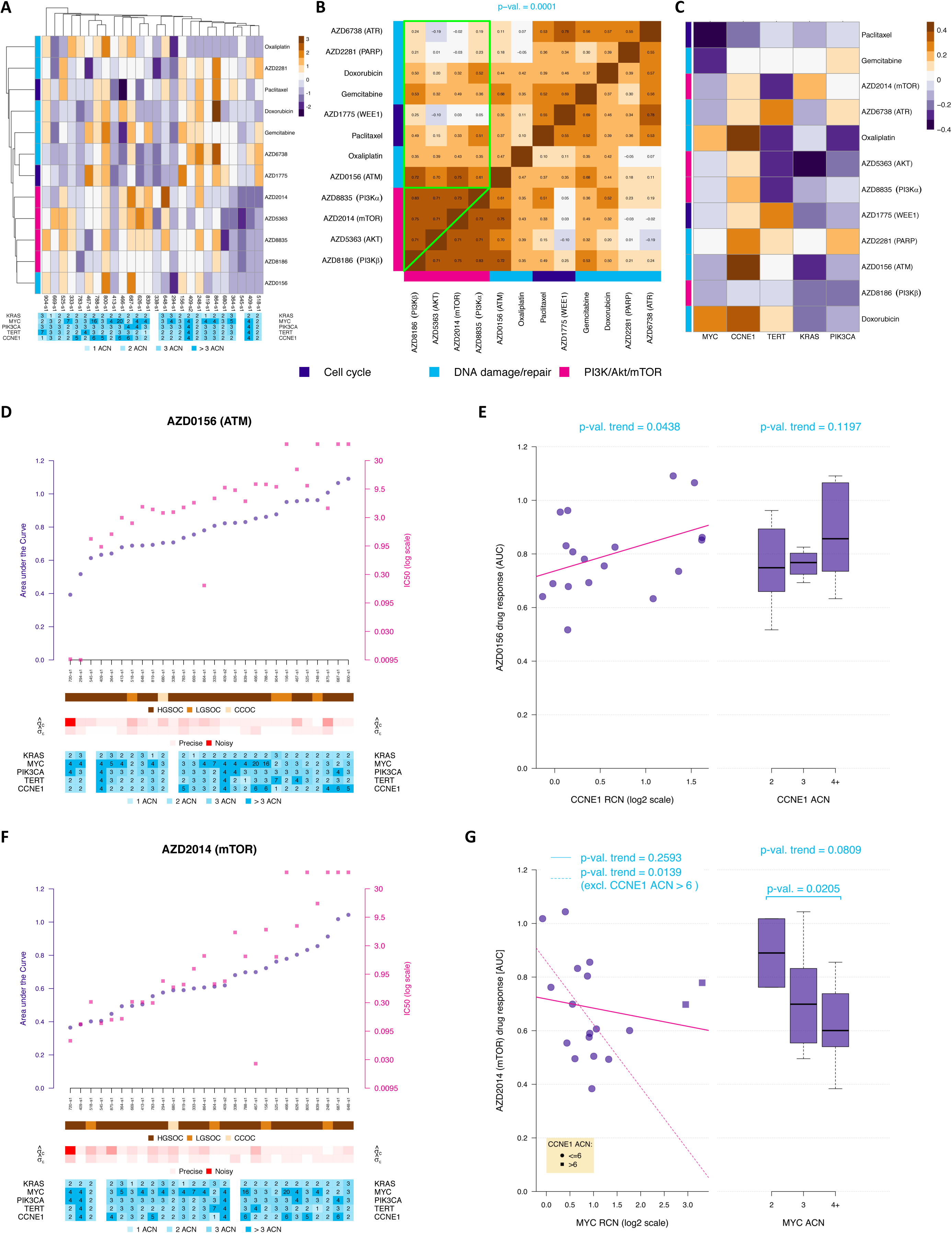
The copy number of important drivers informs response to new targeted therapies. **A**. Heatmap showing how each sample responds to each individual drug. Heatmap colours are obtained from a scale based on Z-score for response. The lower table shows the adjusted copy number for 5 selected genes for each sample. The tricoloured bar on the left shows the pathways affected by each specific drug. **B.** Heatmap showing the Spearman’s Rho correlation between *in vitro* response to different drugs. Response to drugs affecting the PI3K pathway (pink) tend to be similar (high correlation values; green triangle), whilst the correlation between response to PI3K drugs and other drugs is much lower (green square). The p-value corresponding to the non-parametric bootstrap test comparing these two sets of correlations is indicated. **C.** Heatmap showing the correlation between copy number in selected genes and drug response measured by AUC. The tricoloured bar on the left shows the pathways affected by each specific drug. **D.** Scatterplot showing AZD0156 (p-ATM inhibitor) response measured in each sample following the same format as in Figure 4B. Cases with sigma-c or sigma-e above 0.5 were considered too “noisy” and removed from the analysis in the plots 5E and G. **E.** Association between CCNE1 RCN and ACN and AZD0156 *in-vitro* response (as in Figure 4C). **F.** Scatterplot showing AZD2014 (dual mTOR inhibitor) response measured in all samples following the format in Figure 4B. **G.** Association between MYC RCN and ACN and AZD2014 *in-vitro* response (as in panel 4C) when including (solid line) or excluding (dashed line) samples with high CCNE1 ACN.

### MYC-amplified HGSOCs are associated with somatic copy number aberrations in genes from the NF1/KRAS and PI3K/AKT/mTOR pathways and activation of the mTOR pathway

We performed a genome-wide pathway analysis using the TCGA cohort and confirmed that expression of mTOR-related genes was highly correlated with MYC expression (Figures 6A and B). Previous work demonstrated that pathologically elevated expression of MYC expression would have a pro-apoptotic effect (48). We therefore hypothesised that activation of the mTOR pathway mediated the anti-apoptotic survival mechanisms in the context of high MYC and that inhibition of mTOR was lethal through activation of apoptosis. When we compared the genomic chromosomal copy number landscape of HGSOC with increased MYC copy numbers against tumours with diploid MYC, we found that there was an increase in the prevalence of SCNAs, predominantly affecting genes from the mTOR, RAS and anti-apoptotic pathways (Figure 6C). More specifically, the increase in SCNAs affecting PI3K and RAS genes was significantly higher, when compared to other cancer genes (p-value <0.0001). Additionally, amplification of PIK3CA, IGFR1, GAB2, PTK6, KRAS and AKT1/2/3, as well as deletion of NF1 and PTEN (which lead to activation of the RAS and PI3K pathways, respectively) were associated with MYC copy number (Figure 6D). This suggests that co-occurrence of SCNAs affecting these genes and MYC amplification is an evolutionary requirement.

**Figure 6.**
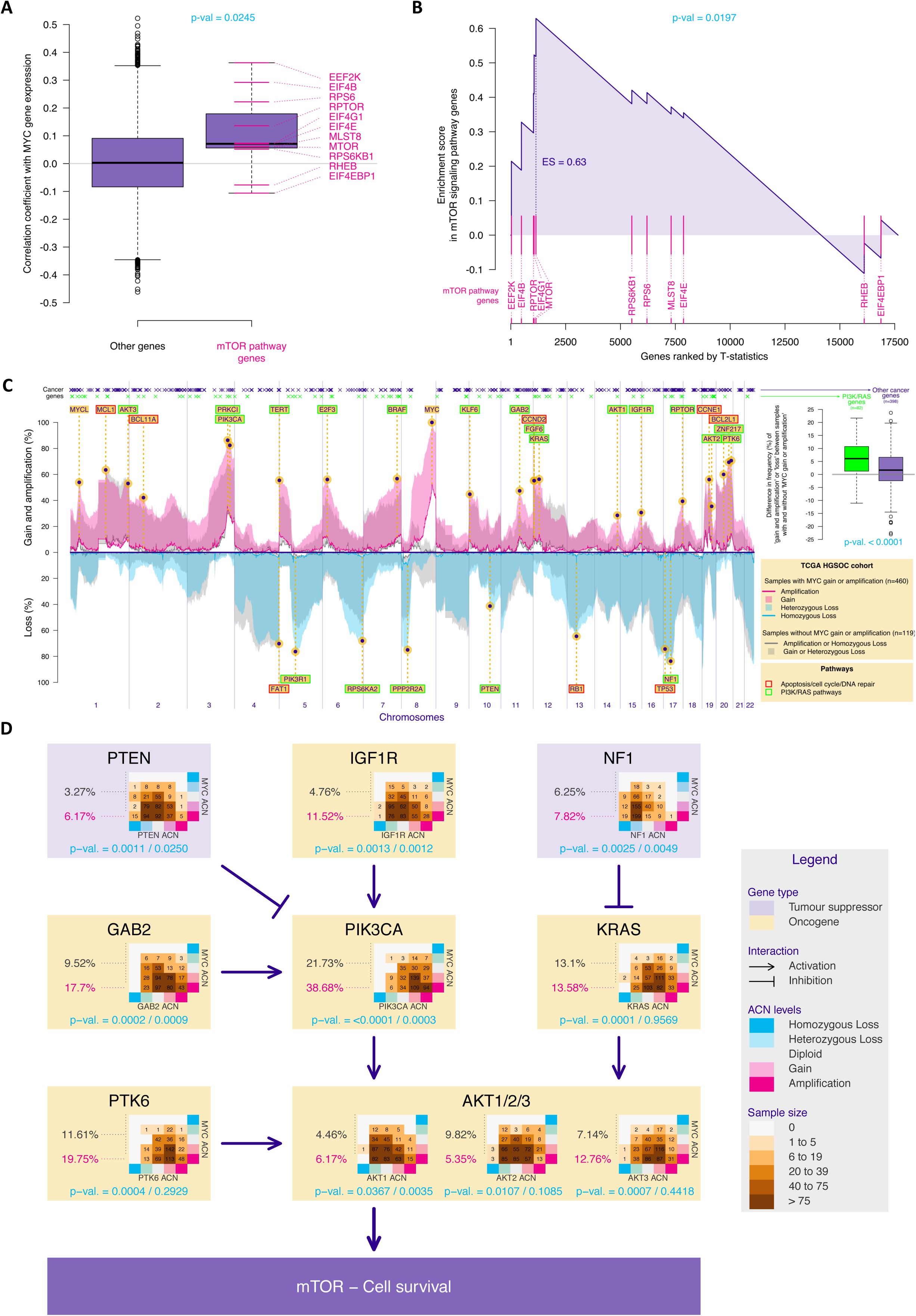
MYC-amplified HGSOCs are associated with SCNAs in genes from the NF1/KRAS and PI3K/AKT/mTOR pathways and activation of the mTOR pathway. **A**. Boxplots showing the Pearson’s correlation coefficient between the gene expression of all genes and the one of MYC, for genes belonging or not belonging to the mTOR signalling pathway. The latter group showed, on average, higher correlation estimates compared to the other group. **B.** GSEA enrichment scores showing enrichment of mTOR signalling pathway genes in Myc-high tumours. The vertical pink lines represent the projection of individual genes from the mTOR pathway onto the gene list ranked by MYC expression level. The curve in blue corresponds to the calculation of the enrichment score (ES) following a standard gene set enrichment analysis (GSEA). The more the blue ES curve is shifted to the upper left of the graph, the more the gene set is enriched in MYC-high genes. The ES score, the normalised ES score (NES) and p-value are also shown in the plot. **C.** Frequency plot showing the distribution of chromosomal amplifications/homozygous losses (solid lines) or gains/heterozygous losses (shaded areas) across the genome in both MYC-amplified/gain (pink for amplifications/gains and blue for losses) and MYC diploid HSOC (gray) in the HGSOC TCGA cohort. The location of a list of functional cancer genes selected in (49) is indicated on top. Cancer genes are colour-coded in green if they belong to the PI3K or RAS pathways based on the Reactome definition. The boxplots (right panel) show, for both PI3K/RAS and other cancer genes, the difference between the frequency of cancer genes SCNAs in tumours with and without MYC amplification or gain. The p-value of the one-sided permutation test of equality of means is indicated. **D.** Diagram showing HGSOC drivers that impact the PI3K pathway and the prevalence of SCNAs across MYC allelic copy numbers (table). For each gene, the p-values corresponding two tests of association between both sets of absolute copy number are indicated (Chi-square test on the left and generalized Cochran-Mantel-Haenszel test for ordered factors on the right) are indicated in turquoise.

### Tumours driven by chromosomal instability share driver SCNAs which are regulated by similar survival pathways

We further explored if HGSOC drivers and survival mechanisms were relevant in the context of other tumours mostly driven by chromosomal instability(50). We compared the genomic landscape of HGSOC (Figure 6C) to the profile of SCNAs in p53-mutant triple-negative breast tumours (TCGA cohort, n=261, Figure 7A; Metabric cohort, n=219, Figure 7B) and squamous lung cancer (TCGA cohort, 501, Figure 7C). We detected an overlap between the genes that are frequently amplified or deleted across these tumours, in the context of MYC amplification or gain. More importantly, we compared the frequency of SCNAs encompassing cancer genes between p53-mutant triple-negative breast tumours or squamous lung tumours, harbouring MYC gain or amplification, with equivalent tumours with diploid MYC. We found a significant increase in the SCNAs encompassing genes from the PI3K and RAS pathways, compared to other cancer genes, in the context of MYC amplification or gain in the TCGA cohorts (TCGA breast cohort: p-value: 0.0001; TCGA lung cohort: p-value < 0.0001). In p53-mutant triple-negative breast tumours from the Metabric cohort, the overall rate of SCNAs was lower, which may have contributed to the absence of similar associations between SCNAs affecting MYC and PI3K/RAS genes (p-value 0.5870). Despite the significant overlap between SCNAs affecting tumours in these three distinct organs, there was a small number of genes that were specifically amplified or deleted in certain tumours (eg. ERBB2 amplification in p53-mutant breast cancers and CRKL amplification or deletion of LRP1B or CDKN2B in squamous lung cancer). Interestingly, SCNAs in the majority of these “private” genes also lead to the activation of the same survival mechanisms, suggesting that they are preserved across different tumours.

**Figure 7.**
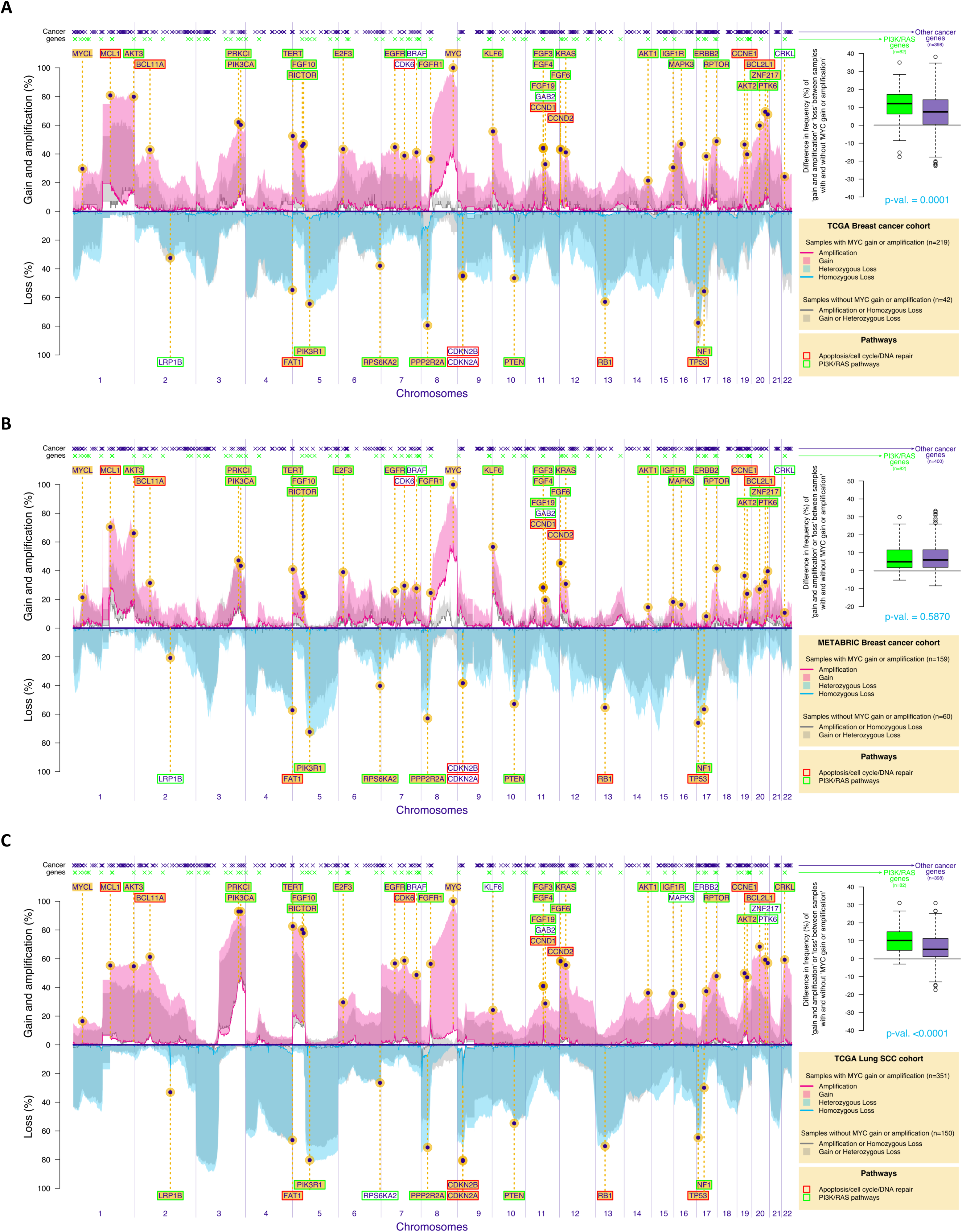
Frequency plots showing the distribution of chromosomal amplifications/homozygous losses (continuous line) or gains/heterozygous losses (shade) across the genome in both MYC-amplified/gain (pink for amplifications/gains and blue for losses) and MYC diploid tumours (gray) in the Breast TCGA cohort (**A.**; triple-negative invasive ductal p53-mutant tumours only), Breast Metabric cohort (**B.**, triple-negative invasive ductal p53-mutant tumours only) and Lung Squamous TCGA cohort (**C.**). The location of a list of functional cancer genes selected in (49) is indicated on top. Cancer genes are colour-coded in green if they belong to the PI3K or RAS pathways based on the Reactome definition. The boxplots (right panel) show, for both PI3K/RAS and other cancer genes, the difference between the frequency of cancer genes SCNAs in tumours with and without MYC amplification or gain. The p-value of the one-sided permutation test of equality of means is indicated.

## Discussion

The extreme genomic complexity of HGSOC has prevented the development of new molecularly targeted therapies. The GISTIC analysis had suggested a number of putative HGSOC drivers but there has been limited functional validation of those(35). We here assumed that CIN-induced SCNAs affecting expression of HGSOC driver genes are positively selected during tumour progression and showed that the correlation between CN and gene expression is higher in frequently amplified HGSOC cancer genes than in non-cancer genes from the same amplicons. Equally, promoters of cancer genes are less methylated than the promoters of non-cancer genes. Overall, this suggests that aberrantly high expression in amplified HGSOC driver genes are crucial for initiation, survival and/or progression of the disease.

Our work showed that the initial clinical drug response in HGSOC can be predicted by high-throughput *in vitro* assays using spheroids, independently of whether samples were collected at diagnosis or after several lines of chemotherapy. This suggests that even after several courses of chemotherapy and development of clinical drug resistance, HGSOC cells can resume their original drug response state when placed *in vitro*, and explains why the predictive value of those assays to assess response in subsequent cycles of chemotherapy was less significant. Previous work suggests that epigenetic silencing of tumour suppressor genes is one of the mechanisms of resistance to platinum and that demethylating agents may revert that resistance (51-53). It is possible that spheroid samples are exposed to a rapid epigenetic reprogramming whist in culture and resume their unmethylated original state (54).

Analysis of the tumorigenic potential of several amplified genes in HGSOC using in vivo models, previously identified GAB2 amplification as a driver of the PI3K pathway activation in HGSOC, that is associated with sensitivity to PI3K inhibition in cell lines (25). Here, we provide evidence that the chromosomal copy number of HGSOC driver genes correlates with the response to specific chemotherapeutic and targeted agents both *in vitro* and in patients. Targeting of early and clonal drivers or of subclonal drivers present in a large proportion of tumour cells should be prioritised (55). We here showed that MYC and PIK3CA gain and amplification were frequent early alterations in HGSOC progression. Deregulated MYC promotes further chromosomal instability by affecting multiple aspects of mitotic chromosome segregation(56, 57). Our *in vitro* results showing an association between MYC copy number and response to paclitaxel were also validated using clinical data where MYC gain or amplification were predictive biomarkers of clinical response to carboplatin and paclitaxel in HGSOC. As increased c-MYC levels have been associated with platinum resistance(58), it is plausible that clinical responses to combined carboplatin and paclitaxel in MYC-amplified tumours are due to paclitaxel effect. Myc-regulated protein synthesis is modulated by the mTOR-dependent phosphorylation of eukaryotic translation initiation factor 4E binding protein-1 (4EBP1), which is required for cancer cell survival in Myc-dependent tumours(59). Therefore, inhibition of mTOR has been shown to be synthetically lethal in MYC-driven haematological tumours (59). Overexpression of c-myc had been shown to induce apoptosis, which could be reverted by overexpression of IGF-1, most likely through activation of the PIK3CA/AKT/mTOR survival pathway(48). Our data showed that MYC amplification or gain co-occur with SCNAs in PI3K genes which induce activation of the mTOR survival mechanisms and that mTOR inhibition was most effective in the context of MYC-amplified HGSOC samples.

By demonstrating that other CIN-driven malignancies (eg. triple negative breast cancer and squamous non-small-cell lung carcinoma) have similar genomic landscapes to HGSOC, our data suggest that survival mechanisms active in HGSOC are present across other CIN-driven tumours. This is corroborated by a recent study analysing the functional evolutionary dependencies in cancer. In this study, co-alteration of PIK3CA and the nuclear factor NFE2L2 (whose transcription is partly regulated by MYC) was a synergistic evolutionary trajectory in squamous cell carcinomas(49). Low-depth whole-genome sequencing is becoming increasingly affordable and available and hence this work supports the rational use of this genomic tools to inform personalised treatments and the design of clinical trials in HGSOC and other CIN-driven malignancies.

## METHODS

### Clinical samples and data

#### Cohort 1

We obtained spheroids with primary tumour cells from ascites of patients with ovarian cancer that were recruited as part of the OVO4 study in Cambridge University Hospitals. Twenty-eight samples from twenty-seven patients were obtained included two matched samples obtained before and after administration of a chemotherapeutic regimen in the same patient.

#### Cohort 2

Solid tumour samples were collected from 64 patients in Cambridge University Hospitals as part of the OV04 study (six of the tumour samples were taken from the same patients that also provided spheroid samples). Samples were handled on ice and processed as soon as possible after surgery. Tumour tissues were fixed for 24 hours in 10% neutral buffered formalin (NBF) before being transferred to 70% ethanol and embedded in paraffin.

#### Clinical data

Clinical data collected included dates of sample collection, surgery, treatment and therapy dates (where applicable), date of death and serial serum CA125-levels from diagnosis. CT reports were obtained and classified into categories which included progressive disease, stable disease, partial response or complete response, according to the RECIST criteria. A summary of the clinical cohort is presented in Table 1 and all the plots summarising clinical data are presented as supplementary material.

### Spheroid isolation

Ascitic fluid was collected from patients with the volume ranging between 100-2000 mL volume. The fluid was centrifuged at 800g for 5 minutes and the majority of the supernatant was removed. Blood clots were removed using a butter muslin cloth and the remaining sample was passed through a 40 µm filter. Spheroids were then recovered after a 10 ml wash with PBS and centrifugation at 1500 rpm for 5 minutes. Next the spheroid fraction was divided in two. One portion of the cell pellet was utilised for DNA extraction. The other portion of the cells were resuspended in filtrated acellular ascitic supernatant and 8% DMSO, transferred to freezing vials and kept in liquid nitrogen. Cells were thawed and kept in media at 37C for 12h before drug screening was performed.

### Ex-vivo drug response/Spheroid assays

An 8-point half-log dilution series of each compound was dispensed into 384 well plates using an Echo® 550 acoustic liquid handler instrument (Labcyte) and kept at -20°C until used. Prior to use plates were span down. 50 µl of organoid suspension were added per well using a Multidrop™ Combi Reagent Dispenser (Thermo-Fisher). Following 5 days of drug incubation, a cell viability assay using 30 µl of CellTiter-Glo® (Promega) was performed. We performed technical triplicates. Due to limited sample availability, biological replicates were only performed in selected samples.

### DNA extraction from FFPE samples

Multiple sections at 10 µm thickness were cut for each FFPE sample depending on tissue size and tumour cellularity. Tumour areas were marked by a pathologist on separate Haematoxylin and Eosin (H&E) stained sections to guide microdissection for DNA extraction. Tumour areas from unstained tissue sections were scraped off and dewaxed in 1 ml of xylene, followed by washing in 100% ethanol. After residual ethanol was evaporated (10 min at 37°C), DNA extraction was performed using the AllPrep DNA/RNA FFPE Kit (Qiagen). DNA was eluted in 40 µl kit ATE elution buffer and quantified using Qubit quantification (Thermofisher, Q32851).

### Genomic profiling

DNA extraction was performed using the DNeasy Blood & Tissue Kit (QIAGEN) according to manufacturer’s instructions. DNA samples were diluted to 75 ng in 15 µl of PCR certified water for subsequent shearing by sonication using the LE220-plus Focused-Ultrasonicator (Covaris) for 120 s at RT (duty factor 30%; Peak incident power 180 W; 50 cycles per burst) with a target of 200-250 bp. Library preparation was subsequently carried out using the SMARTer Thruplex DNA-Seq kit (Takara) with each sample undergoing 7 PCR cycles for library amplification and sample indexing. sWGS libraries were cleaned using AMPure beads, according to the manufacturer’s recommendations, and eluted in 20 µl TE buffer. Quality and quantity of sWGS libraries were assessed using a D5000 genomic DNA ScreenTape (Agilent, 5067-5588) on the 4200 TapeStation System (Agilent, G2991AA). Libraries were pooled and sequenced using the Paired-End 50 mode and S1 flowcell on the NovaSeq. Genomes were aligned to the GRCh37 reference genome and relative copy number data was obtained using the qDNAseq package(24).

### Statistical analysis

All statistical tests were two sided, unless stated otherwise.

#### a Test of equality of the location parameter between two populations

In the Figures **2B, 2C** and **6A**, we used Wilcoxon rank sum non-parametric tests (also known as Mann– Whitney-Wilcoxon tests) with continuity correction (*wilcox.test* function of the *stats* R package) to assess if the location shift between two populations is different from 0. Similar conclusions were obtained when considering Welch t-tests (*t.test* function of the *stats* R package) and median test (*median_test* function of the *coin* R package). In the Figures **6C**, **7A, 7B** and **7C**, we used one-sided permutation tests (considering R=10’000 resampling of the TCGA or METABRIC patients) to assess if the mean of the differences (in %) of ‘amplification or gain’ or ‘loss’ (reverse scale) levels of tumours with or without MYC gain or amplification are different for cancer genes (49) belonging or not to the Reactome PI3K or RAS pathways. Triple-negative breast tumours from TCGA cohort (Figure **7A**) were selected based on the Code 8500/3 from the International Classification of Diseases for Oncology, Third Edition ICD-O-3 Histology Code. Triple-negative breast tumours from Metabric cohort (Figure **7B**) were selected based on negativity for ER, PR and HER2 status.

#### b Estimates, test and representation of the association between paired samples

In the Figure **2B**, we estimated the Pearson’s product moment correlation coefficient (*cor* function of the stats R package) between normalised gene expression (obtained with the function *cpm* of the *edgeR* R Bioconductor package) and respective chromosomal copy number for each gene in the HGSOC TCGA database. Similar conclusions were obtained when considering Spearman rank-based correlations.

In the Figure **2D**, we used the Pearson’s product moment correlation coefficient test (function *cor.test* of the *stats* R package), to assess if the level of association between 2 variables is different from 0. In the Figure **2D**, the covariance/correlation matrix and the vector of means were normalised (by considering the quantiles of a standard normal distribution corresponding to the observed probability point of each gene) since they are the sufficient statistics of the bivariate normal distribution. Representation of the level of association between the normalised variables of interest was then obtained by displaying ellipses corresponding to the quantile 0.95 of the bivariate normal distributions of interest. In the Figure **3D**, we used Spearman’s rank-based statistics (*cor* function of the *stats* R package) to describe the level of association between paired AUC and IC50 since those measures are not bivariate normal.

In the Figure **5B**, the association between the *ex-vivo* response to paired drugs was defined by means of Spearman rank-based correlation (cor function of the stats R package). We estimated (i) the level of association between ex-vivo response to each pair of drugs affecting the PI3K/Akt/mTOR pathway, (ii) the level of association considering all paired drugs, including one drug from this PI3K/Akt/mTOR group and one from another group, and compared their mean by means of a non-parametric bootstrap test considering 25000 samples (the corresponding p-value is indicated).

In the Figure **6A**, we estimated the Pearson’s product moment correlation coefficient (function *cor* of the *stats* R package) between expression of MYC and expression of all the other genes from the HGSOC TCGA cohort.

#### c Response to treatment estimates

in Figures **3C** and Supplementary Figures 3A-Z, response to treatment was estimated using area under the curve (AUC) and IC50. Drug response measures were standardised by dividing the original values by the median drug response observed in the control group of each drug and sample. This standardised drug response measures were then modelled as a function of the dose (on the log scale) by means of a 4th degree polynomial robust regression, fitted by means of the function *lmrob* of the R package *robustbase*. Drug response measures that obtained a robust weights smaller than 0.4 (out of a range which spreads from 0 for outliers to 1 for non-outliers) were considered as outliers. After excluding outliers, we modelled the standardised drug response measures as a function of the dose (on the log scale) by means of M-splines. AUC and IC50 were estimated using the I-splines (which correspond to the integrals of M-splines). The alternative use of a five-parameter log-logistic fit (*drm* function of the *drc* R package with function LL2.5) led to similar AUC and IC50 estimates.

In the Figures **4B, 4D**, **5D** and **5F**, the control and epsilon sigma estimates represented in levels of red respectively correspond to (i) the observed standard deviation of the (standardised) drug responses of the control group per sample and to (ii) the standard deviation of the difference between the (standardised) observed drug respond and corresponding M-spline fits per drug.

#### d Linear relationship between continuous variables with independent samples

In the left plots from the Figures **4C, 4E**, **5E** and **5G**, we used linear regressions, fitted by means of the function lm of the stats R package, to model the relationship between drug response (AUC) and MYC relative copy number (on the log scale). The p-values of the one-sided Wald t-test corresponding to the slope parameter of each fit are indicated.

In the left plot of the Figure **5G**, the trend between drug response and MYC relative copy number was also fitted when excluding the two observations corresponding to samples with high CCNE1 amplification (ACN > 6; patients 466 and 788). The direction of the relationship was pre-specified(40).

We only considered the first sample for patient 409, since the linear model requires independent observations. Equally, the samples from patients 720 and 875 were excluded from these analyses due to extreme variability in the results demonstrated by the high standard deviations in the (standardised) drug responses of the control group (Supplementary Figures 3A-Z).

#### e Trend test between an ordinal and a continuous variable

In the Figure **3B**, we used an ordinal regression (*clm* function of the *ordinal* R package) as well as Jonckheere-Terpstra test (*jonckheere.test* function of the *clinfun* R package) to assess the level of association between the ordinal CT responses and the difference in CA125 on the log2 scale before and after treatment. Both methods lead to a p-value < 2.2e-16.

In the right plots of Figures **4C, 4E**, **5E** and **5G**, we used one-sided Jonckheere-Terpstra tests (*jonckheere.test* function of the R package *clinfun*), to test for ordered differences in drug response between the 3-level of MYC absolute copy number variable (2, 3 and 4+), assuming the same trend direction as in the plots on the left.

In the Figures **4E** and **5G**, we used Student’s t-tests, (*t.test* function of the stats R package), to investigate a difference in means between two levels of KRAS and MYC ACN. The second sample of patient 409 and the samples of patients 720 and 875 were excluded from these analyses for the same reasons described above.

#### f Linear relationship between continuous variables with dependent samples

As patients typically have several lines of treatment and as their responses to treatment are likely not independent, linear mixed models with patients as random effects (to take the within-patient dependence into account) were used in analyses related to the clinical drug response. Models were fitted by means of restricted maximum likelihood (REML; *lme* function of the *nlme* R package).

In the Figure **3E**, a random intercept model was fitted for each drug and the p-values of the two-sided Wald t-test corresponding to the slope parameter was indicated in the legend. A linear regression was preferred when the subset of data corresponding to a drug were independent (only one observation per patient).

In the Figure **3F**, an heteroscedastic mixed linear model with [i] patients as random effects, [ii] AUC, line number (as a three-level factor: 1, 2 and 3+) and the interaction between AUC and line number as fixed effects and [iii] residuals as a power function of the duration of the chemotherapy cycle (to account for the fact that the variability increases with duration) was fitted. The table on the bottom right of Figure F3F shows the p-values obtained when comparing the slope of each line number (rows) to 0 (no relationship between ex-vivo and clinical response) and other line numbers (pairwise comparisons) using a multiplicity correction taking the dependence between these tests into account, as implemented in the function *ghlt* of the *multcomp* R package. Note that the same conclusions were obtained when fitting alternative models (e.g. when not considering heteroscedasticity residuals). In the Figure **4F**, random intercept models, considering patients as random effects and MYC relative copy number (on the log scale) as the fixed effect were fitted with or without controlling for tumour purity (obtained from TP53 mutant allele frequency). The p-values indicated correspond to the Wald t-test of the slope parameter obtained with or without tumour purity in the model.

#### g Copy number frequency plots

The copy number frequency plots in the Figures **2A**, **6C**, **7A, 7B** and **7C** were obtained by computing the percentage of [i] homozygous loss, [ii] heterozygous loss, [iii] gain and [iv] amplification for the subset of samples of interest (all TCGA or spheroids samples in **2A**; for samples showing MYC gain or amplification or neither of both in **6C, 7A, 7B** and **7C**). Amplification (homozygous loss) and combined gain and amplification (homozygous and heterozygous losses) were displayed for each gene and subset of interest. Considering a log2 relationship between relative and adjusted copy number, we used adjusted copy numbers of 0.5, 1.5, 2.5 and 4.5 (Figure **4A**) to define the limits of the subgroups in the spheroid cohort from Figure **2A**. in the Figures **2A**, **6C**, **7A, 7B** and **7C**, we considered the absolute copy number available for the TCGA and METABRIC.

#### h Differential gene expression analysis

In the Gene Set Enrichment Analysis (Figure **6B**, we obtained a list of ranked genes according to the t-statistic corresponding to the change in mean gene expression intensities between HGSOC TCGA samples with MYC-low and MYC-high (lower and upper quartile of MYC expression respectively). We discarded 6522 out of 24165 genes with low counts across all samples (counts-per-million below 0.5 in more than 90% of the samples; from the cpm function of the edgeR R Bioconductor package). Differential expression analysis was performed using the voom function of the *edgeR* R Bioconductor package (with default options).

#### i Gene score enrichment analysis

To assessed whether the up-regulated genes in MYC-high compared to MYC-low tumours were enriched for mTOR pathway genes, we obtained a list of genes belonging to the mTOR signalling pathway from the Bioplanet database (https://tripod.nih.gov/bioplanet/, the pathway “Mammalian target of rapamycin complex 1 (mTORC1)-mediated signalling”) and ran a gene set enrichment analysis (*fgsea* function of the R Bioconductor package *fgsea*; Figure **6B**), using the mTOR gene list and the list of ranked genes of the differential gene expression analysis described above, to obtain the enrichment score estimate and corresponding p-value.

#### j Clustering analyses

In the Figures **5A**,**5B** and **5C**, we used hierarchical clustering to group samples, drugs or genes. In the Figure **F5A** and **F5C**, samples, drugs and genes were grouped according to a ‘complete’ hierarchical clustering based on euclidian distances (*pheatmap* function of the *pheatmap* R package and *hclust* function of the *stats* R package). In the Figure **5B**, the drugs were grouped according to a ‘complete’ hierarchical clustering based on euclidian distances defined on the drug Pearson’s correlation matrix (*hclust* function of the *stats* R package).

#### k Tests of association of categorical or ordered factors

In the Figure **6D**, we present the association between the absolute copy number (ACN) of selected genes and the ACN of MYC in two different way. Firstly, we considered the ACN vectors as non-ordered factors and used the Chi-square tests (*chisq.test* function of the stats R package). Secondly, we considered them as ordered factors and used the generalized Cochran-Mantel-Haenszel tests of association of ordered factors (*CMHtest* function of the *vcdExtra* R package).

## ACKNOWLEDGEMENTS

We are very grateful for Prof. Samuel Aparicio’s comments and careful review of the manuscript and Dr. Suzanne Carreira’s review of mutational data in a previous clinical trial.

## CONTRIBUTIONS

**F.C.M.** developed the concept and directly supervised the study; **F.C.M., R.C., C.S.** and **J.D.B.** obtained funding for the study; **F.C.M., D.S.** and **M.V.**, designed and performed the drug screens; **F.C.M., A.A.** and **K.H.** collected the clinical data, **F.C.M.** and **D.L.C.** generated hypotheses, analysed and interpreted molecular, experimental and clinical data, wrote the initial manuscript draft and edited the final version; **D.L.C.** led the bioinformatic and statistical analysis of the clinical, experimental and genomic data and designed the manuscript figures **I.S.** performed the TCGA SCNA/gene-expression and gene-enrichment analysis and provided considerable bioinformatic support throughout the work. **C.M.S., M.V.,A.P.,J.H.** collected and processed primary tumour and spheroid samples or prepared their SGWS libraries. **M.A.** performed the clonality analysis using the multi-regional HGSOC sequencing data**. S.C., L.C., B.D.** facilitated access to AZ drugs and drug screen facilities, and assisted the analysis of results from the drug screen; **G.F., H.B., K.H., J.L., P.B., R.C., B.B.**, helped with the analysis by assisting in data collection or establishing avenues of enquiry related to patient clinical and molecular characteristics. **T.B.K.W., M.E., N.M., K.L.** provided guidance on the bioinformatic analysis. **S.S.** provided processed multi-regional HGSOC sequencing data. **B.B., S.S., I.M., C.C., G.E., C.S., J.D.B.**, provided valuable guidance in the analysis and/or revision of the data. **C.S.** developed the concept and supervised the evolutionary multi-regional analysis. **J.D.B.** is the lead for the OVO4 study and supervised the genomic analysis. **C.S.** and **J.D.B.** hosted and provided supervision to **F.C.M.** throughout the study, reviewed the full analysis and helped preparing the final manuscript. All authors reviewed, provided valuable contributions and approved the final submitted manuscript.

---

**Supplementary Figure 1.**
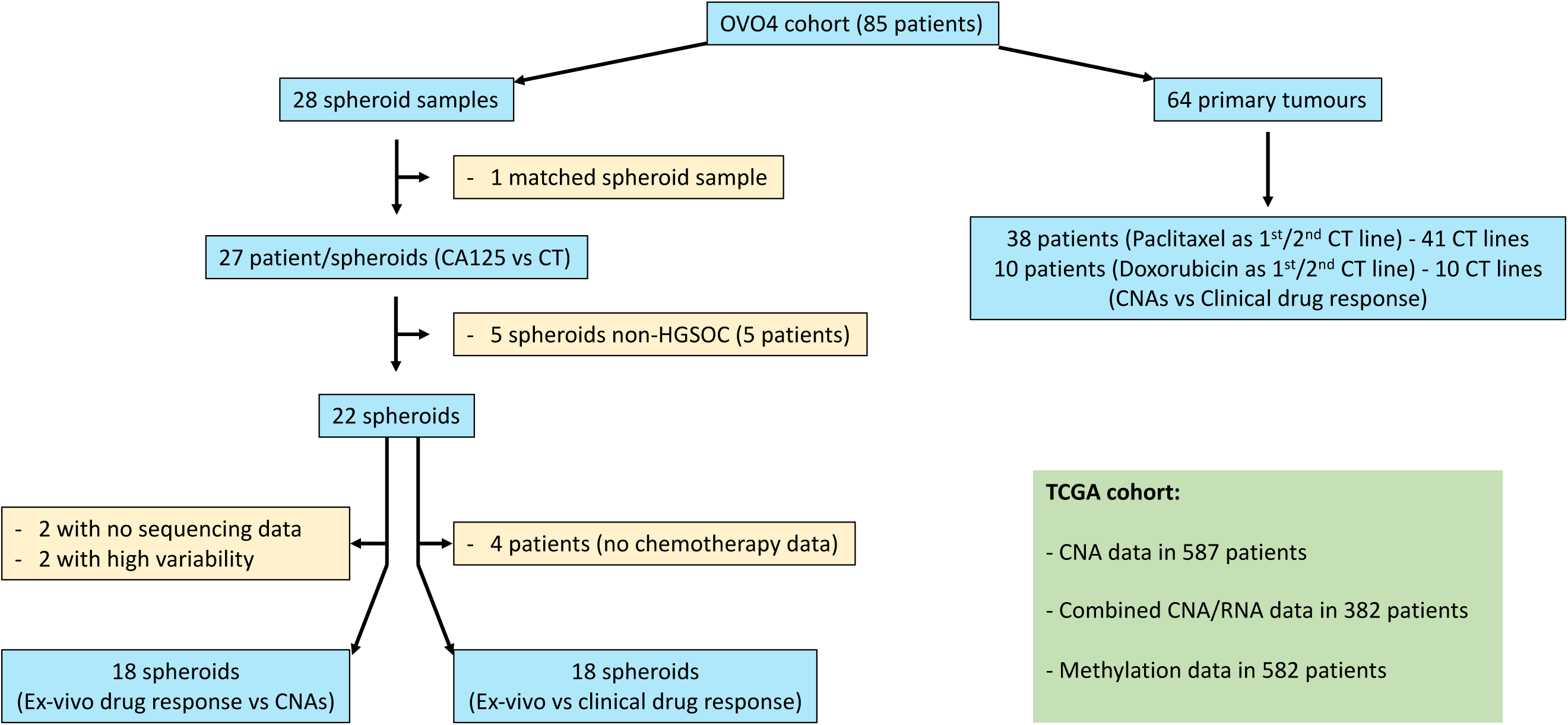

**Supplementary Figure 2.**
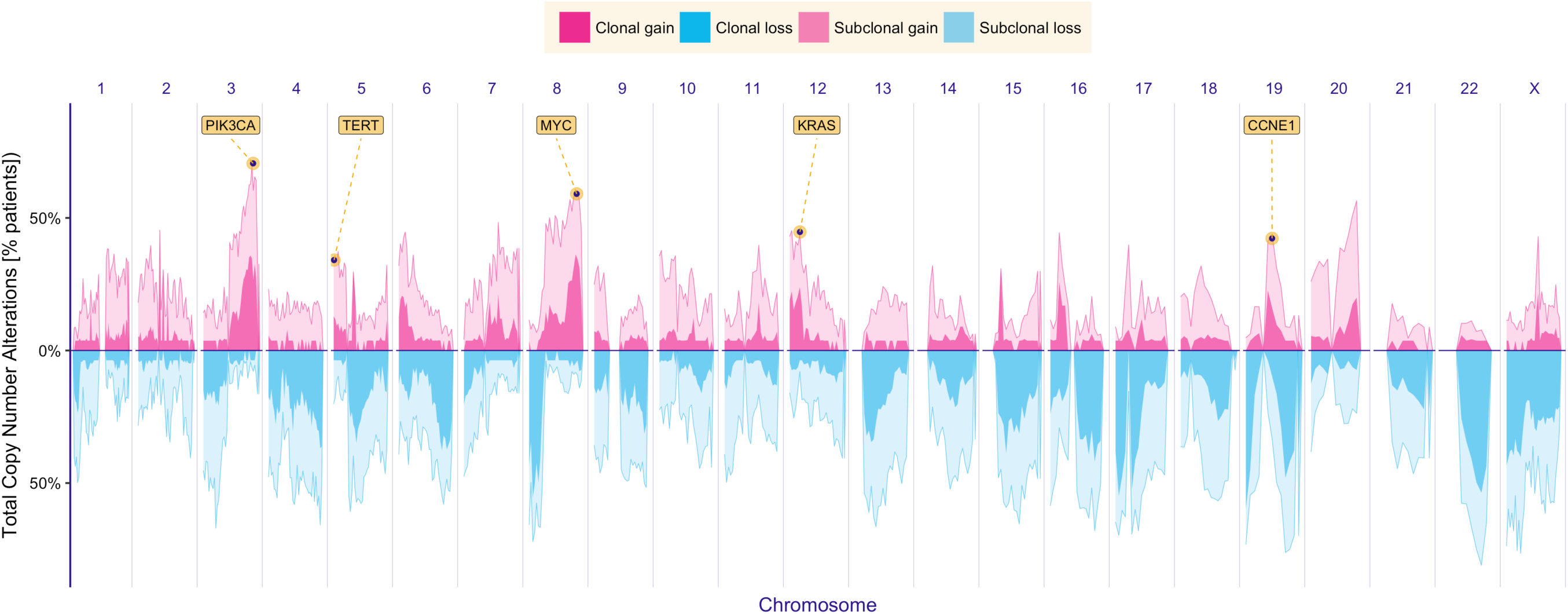

**Supplementary Figure 4.**
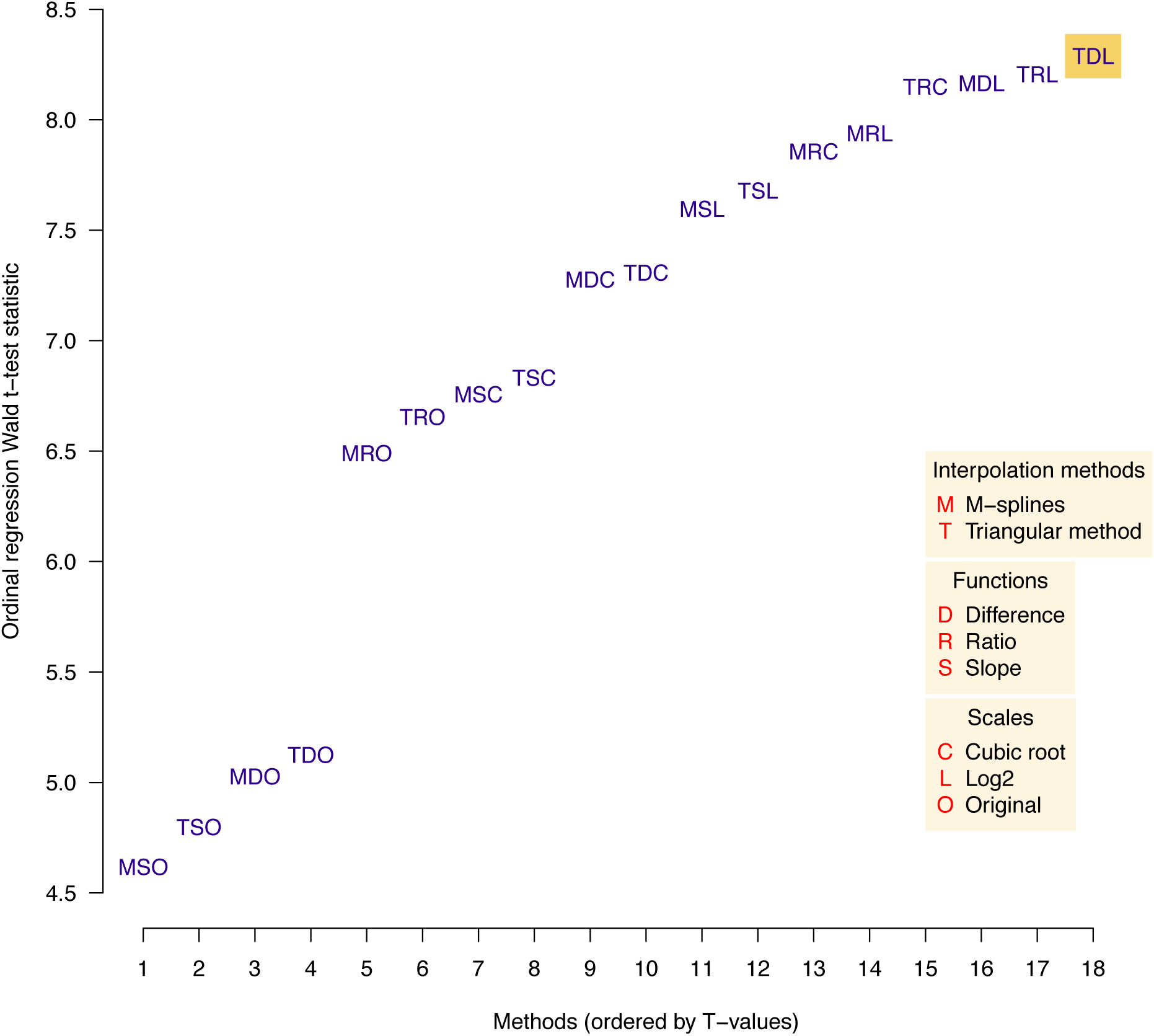

## Notes

### Competing Interest Statement

The authors have declared no competing interest.

