## Supplementary Figures 3A-Z for "Somatic chromosomal number alterations affecting driver genes inform *in-vitro* and clinical drug response in high-grade serous ovarian cancer": SupplementalFigures3A-Z.pdf

### Suppl. Figure 3A -- Patient 156 -- LGSOC -- Stage III

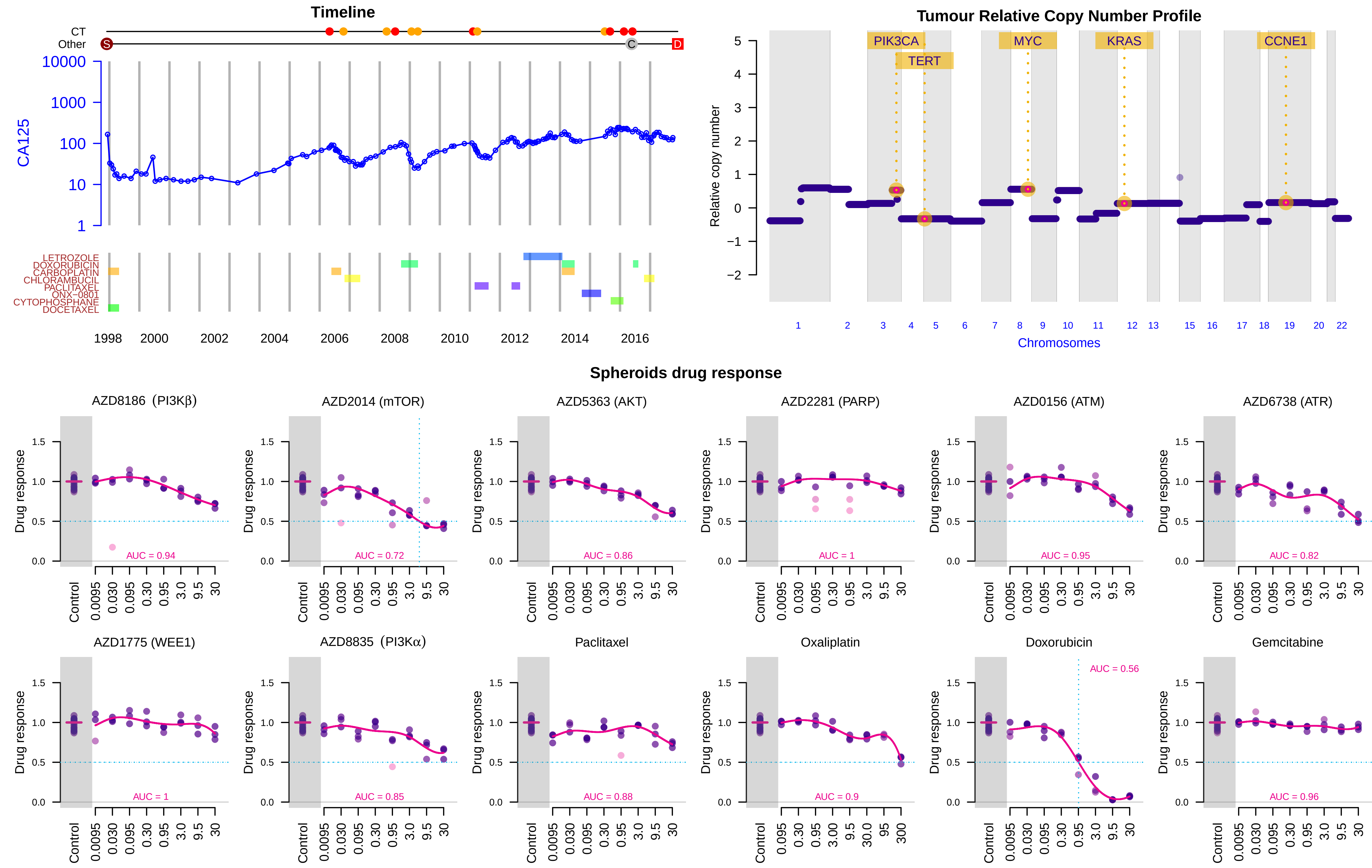

### Suppl. Figure 3B -- Patient 248 -- LGSOC -- Stage III

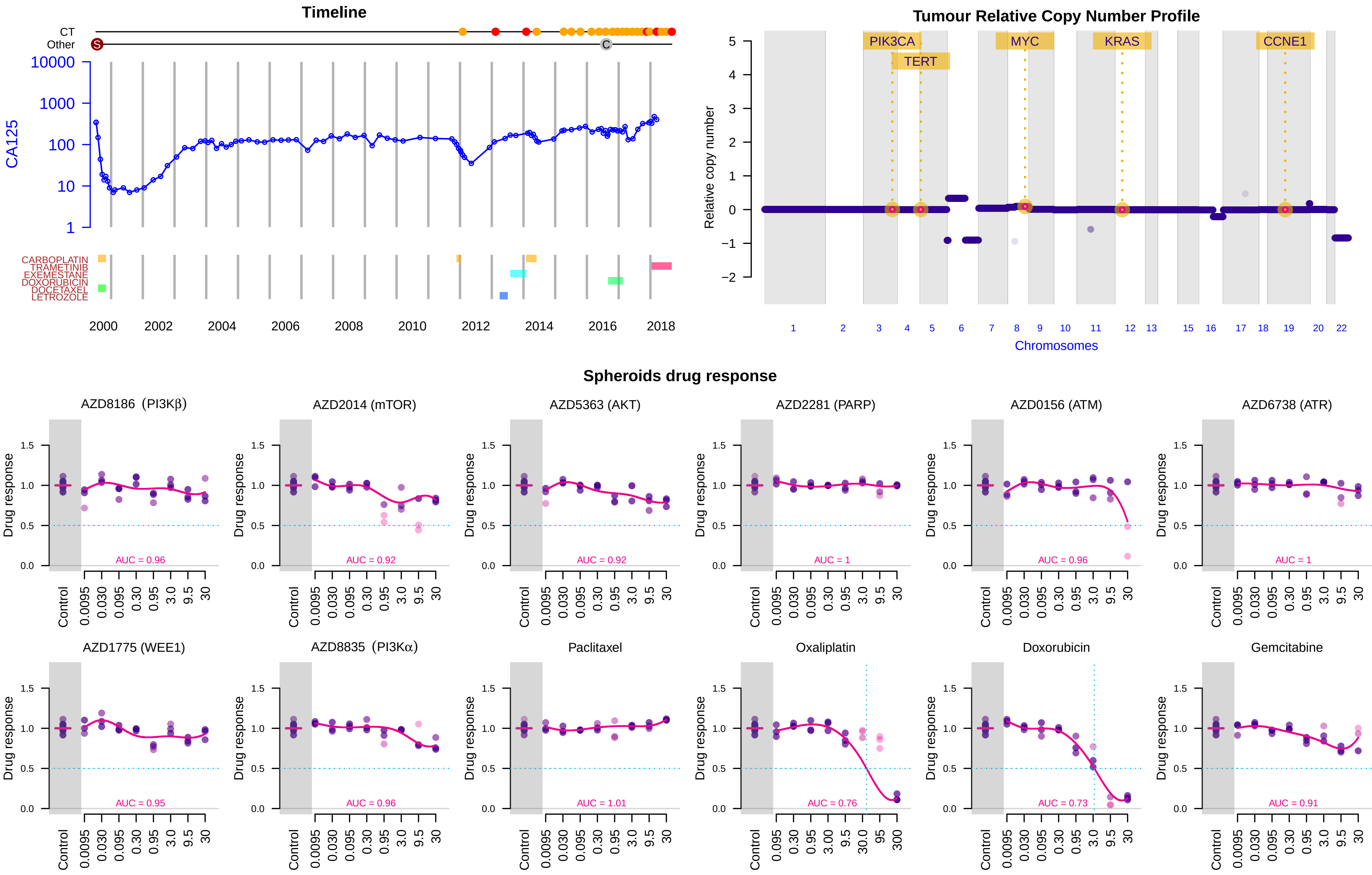

### Suppl. Figure 3C -- Patient 294 -- HGSOC -- Stage III

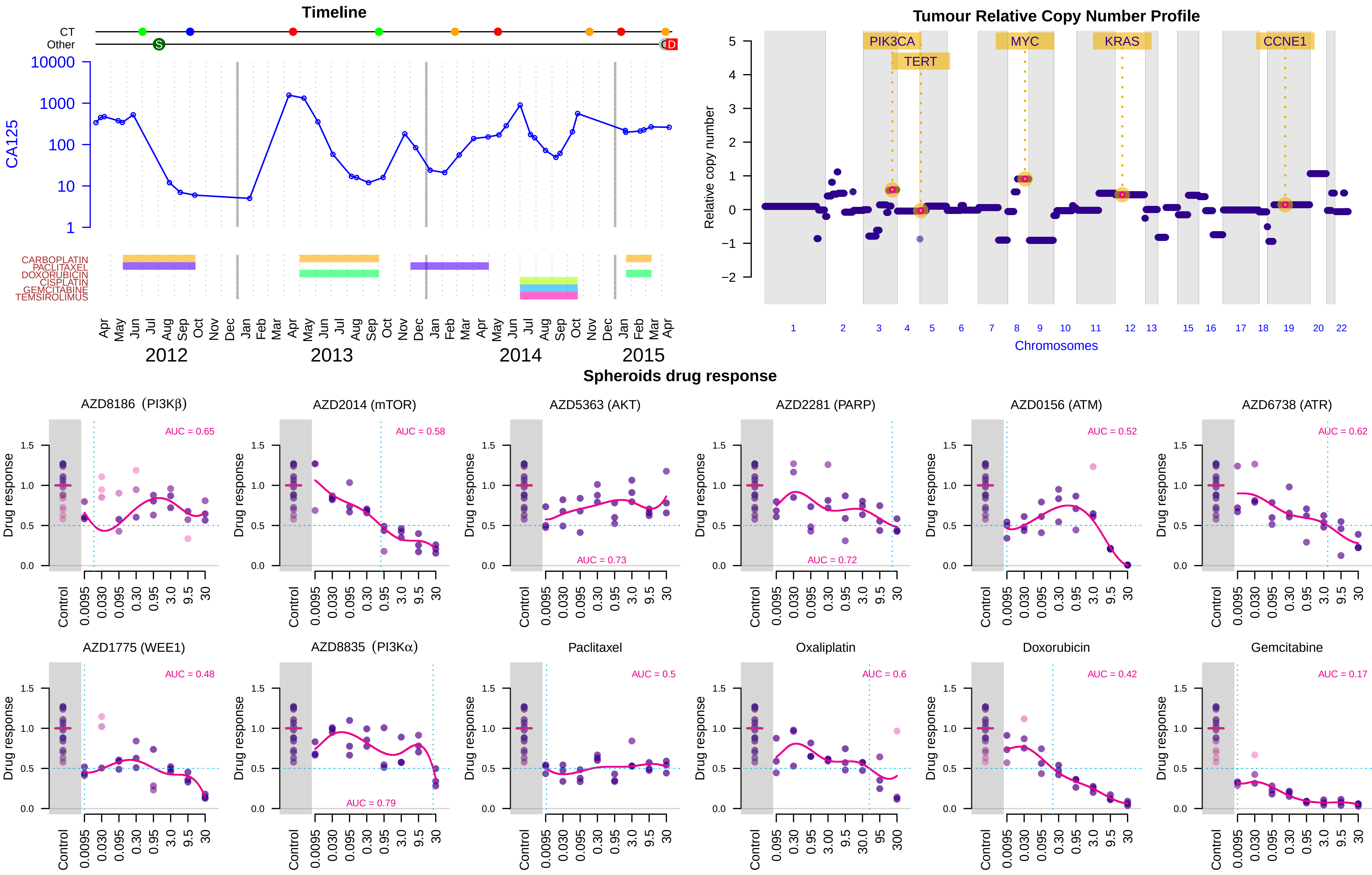

Suppl. Figure 3D -- Patient 333 -- HGSOC -- Stage IV

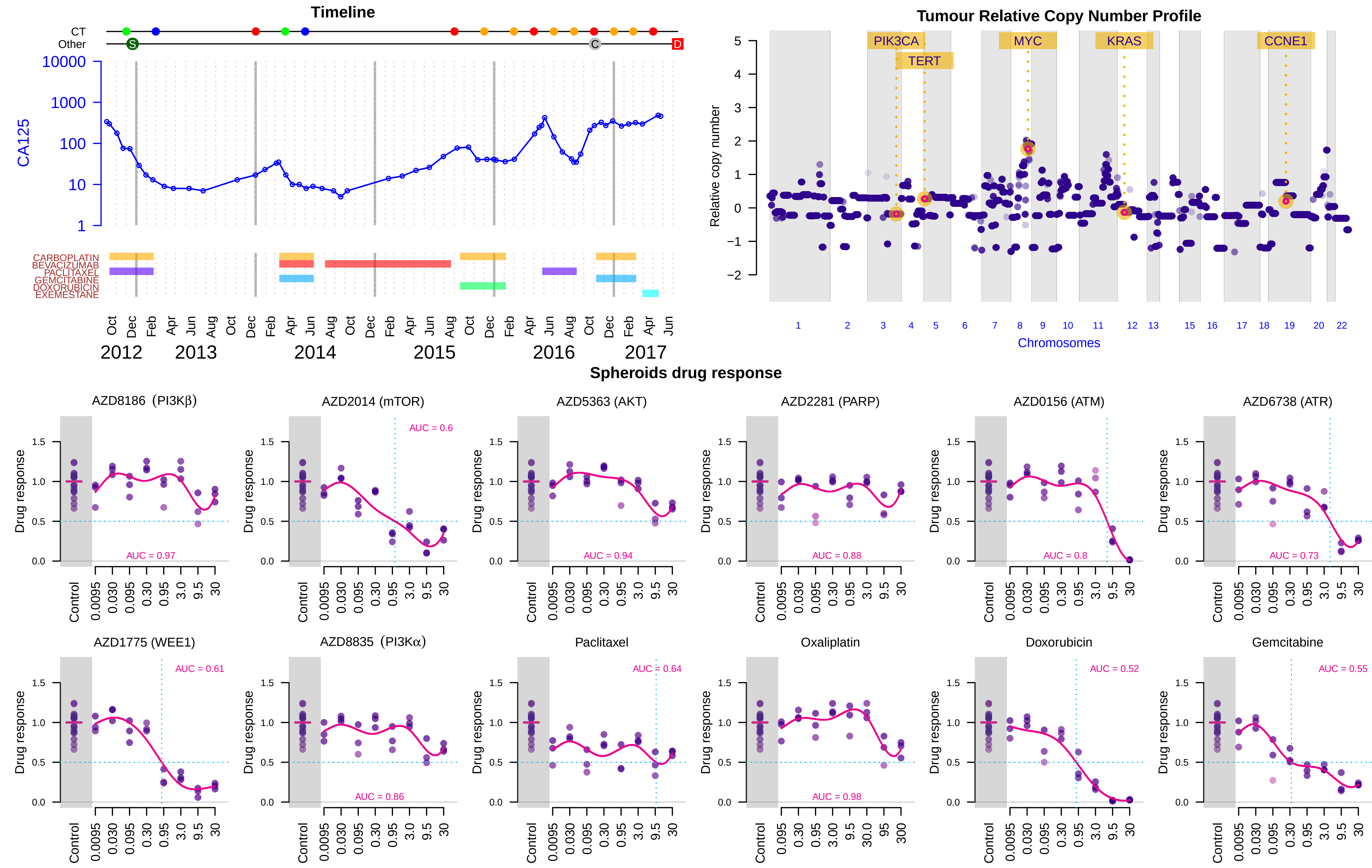

Suppl. Figure 3E -- Patient 364 -- HGSOC -- Stage IV

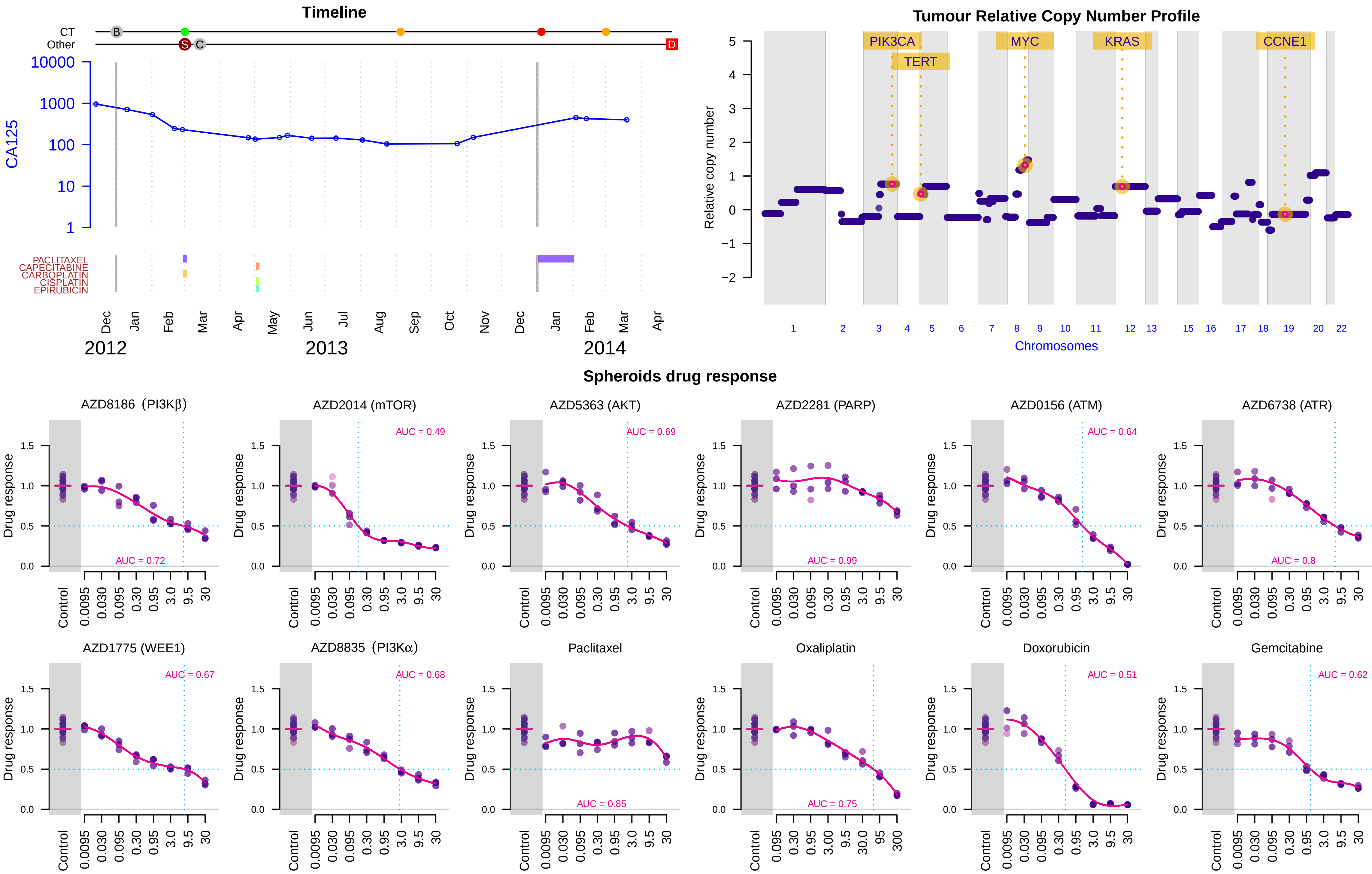

### Suppl. Figure 3F -- Patient 409 (First sample) -- HGSOC -- Stage IV

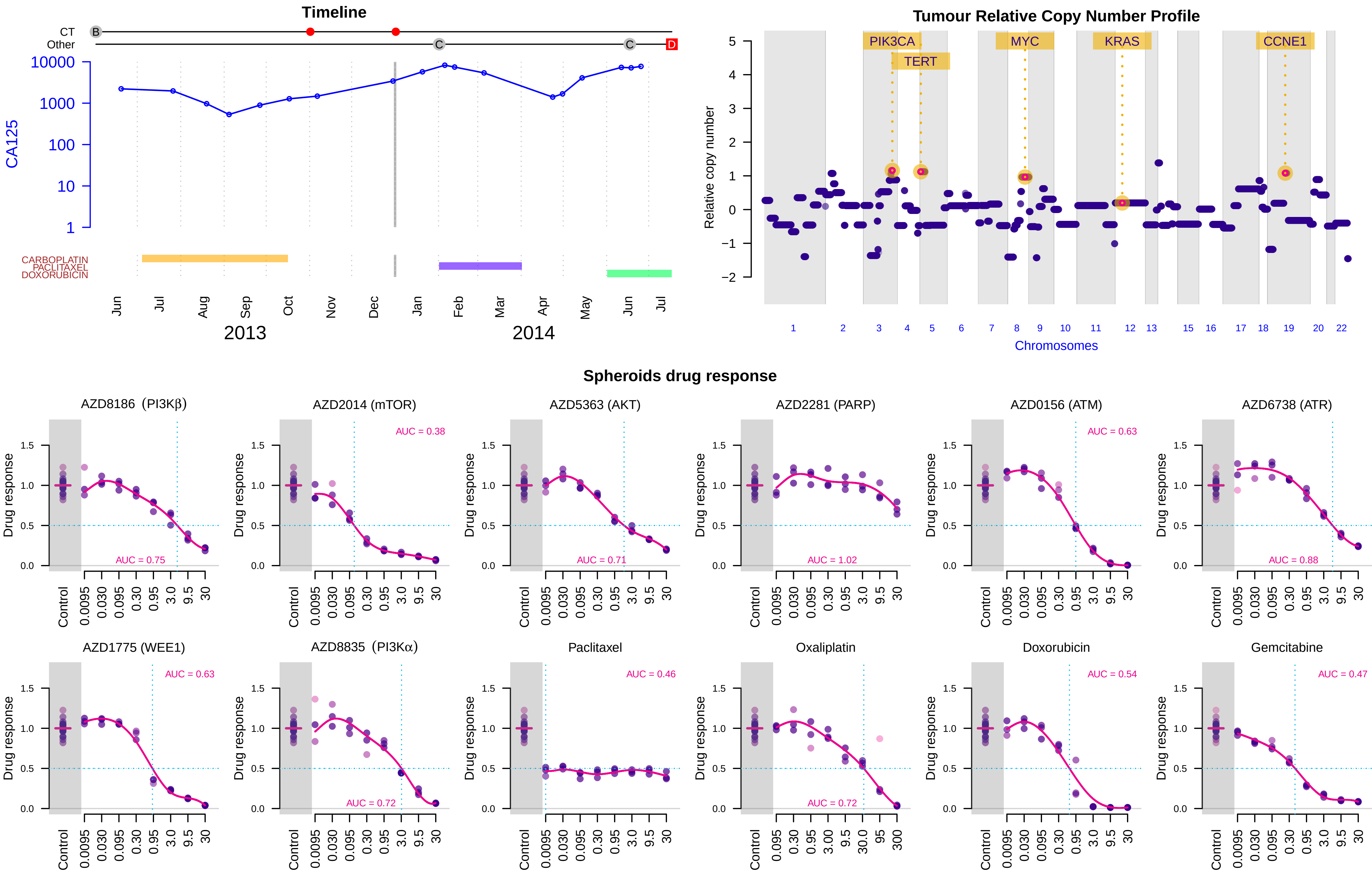

Suppl. Figure 3G -- Patient 409 (Second sample) -- HGSOC -- Stage IV

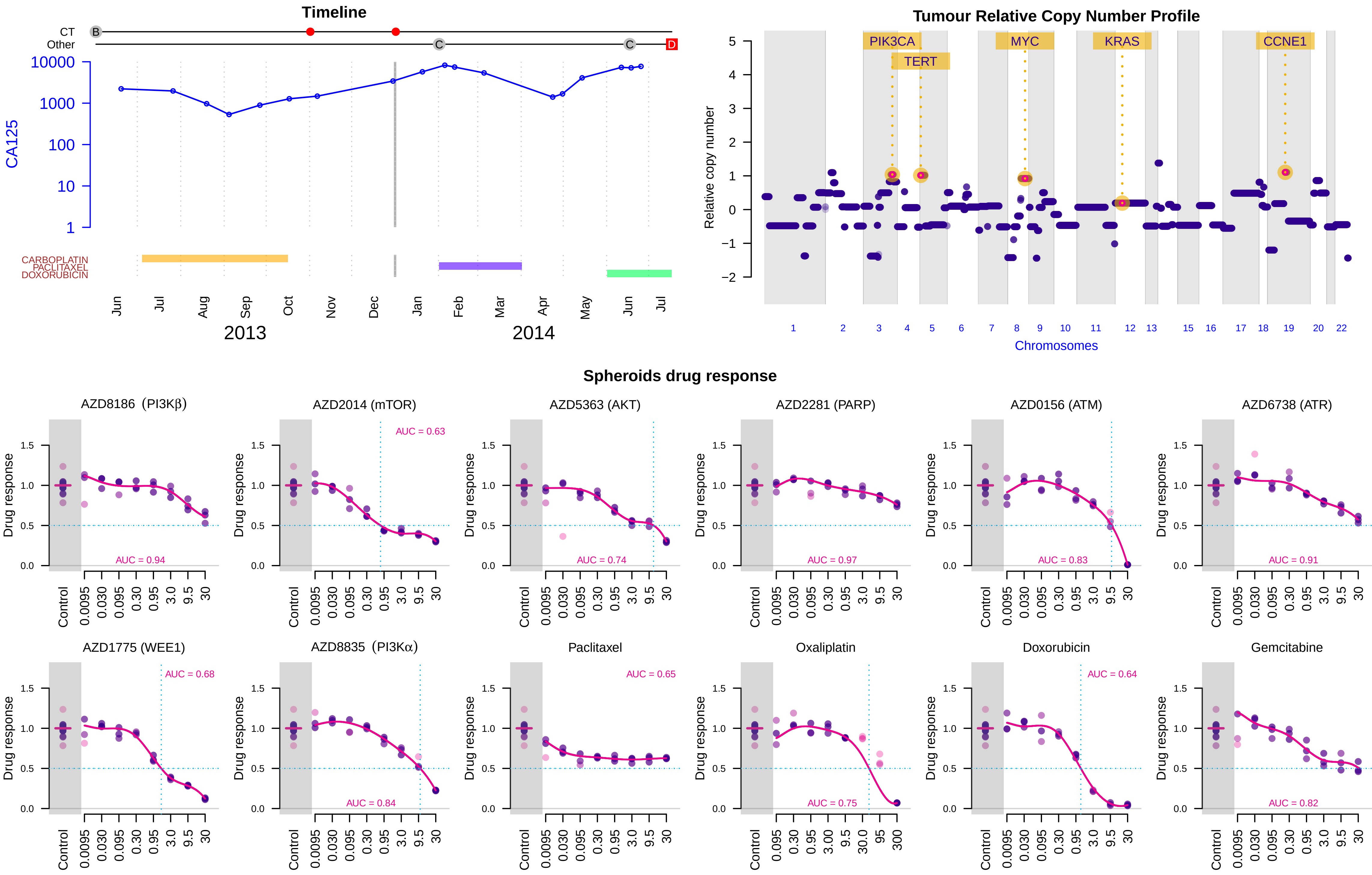

Suppl. Figure 3H -- Patient 413 -- HGSOC -- Stage III

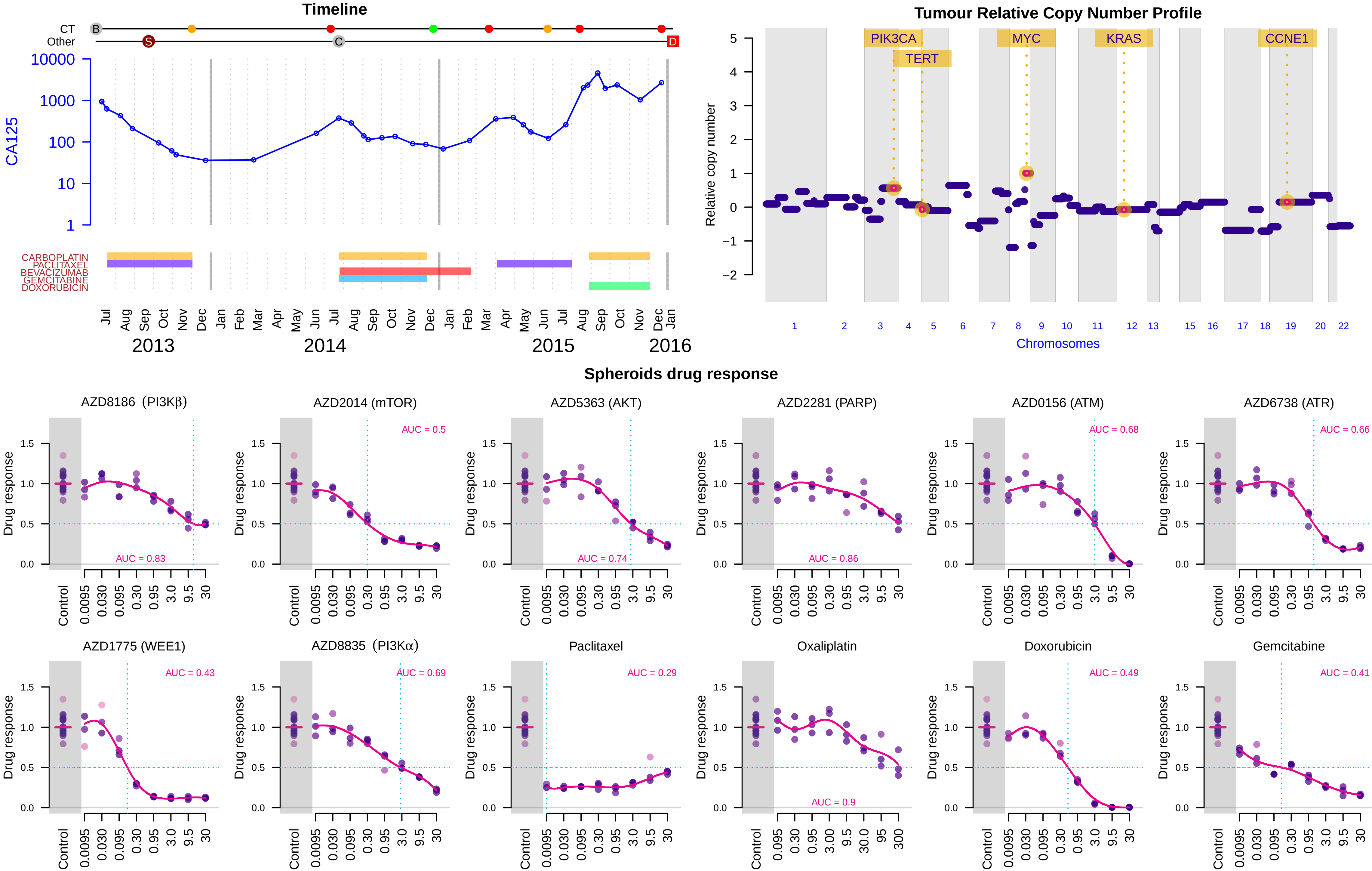

Suppl. Figure 3I -- Patient 466 -- HGSOC -- Stage III

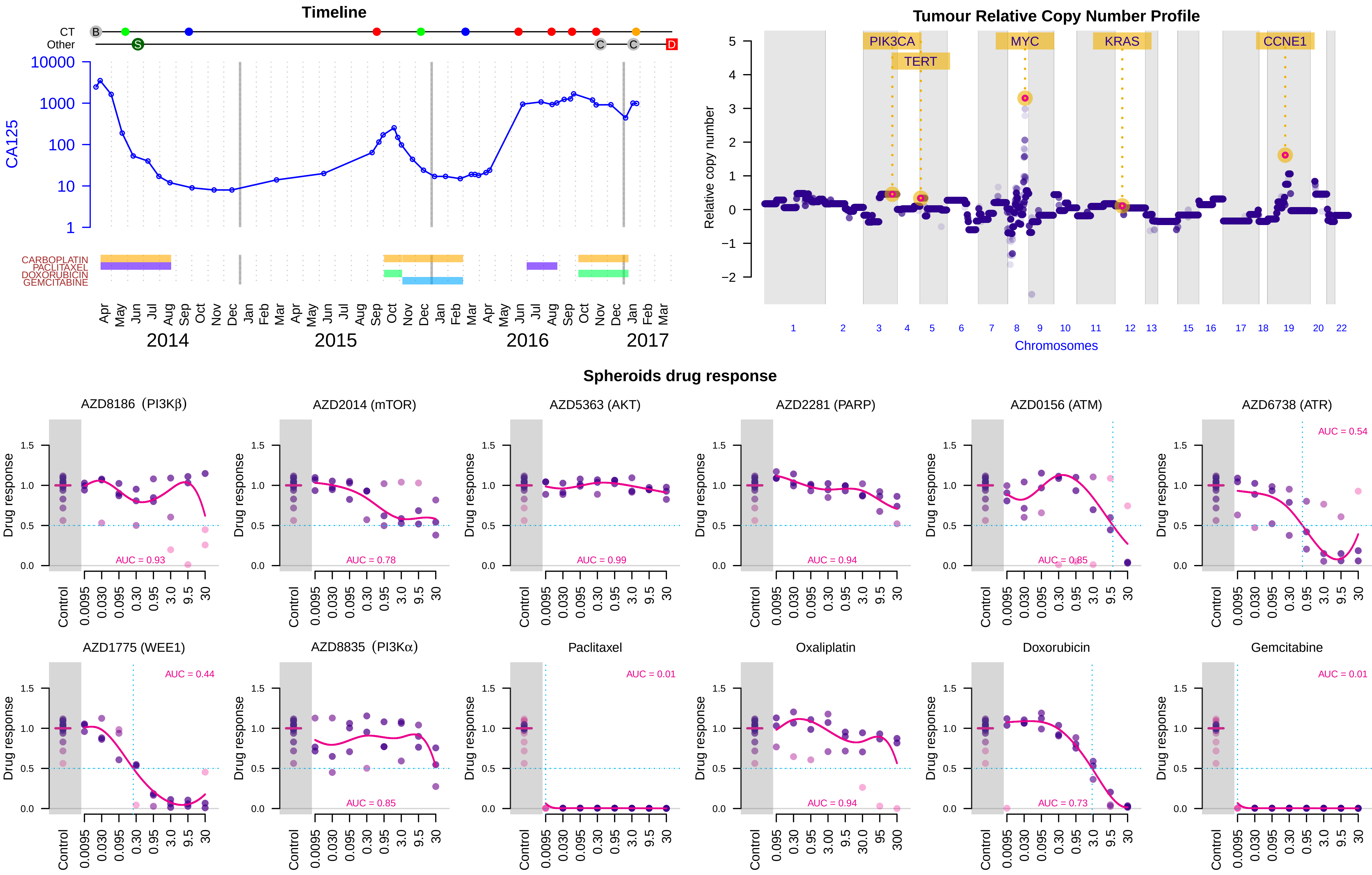

### Suppl. Figure 3J -- Patient 467 -- HGSOC -- Stage III

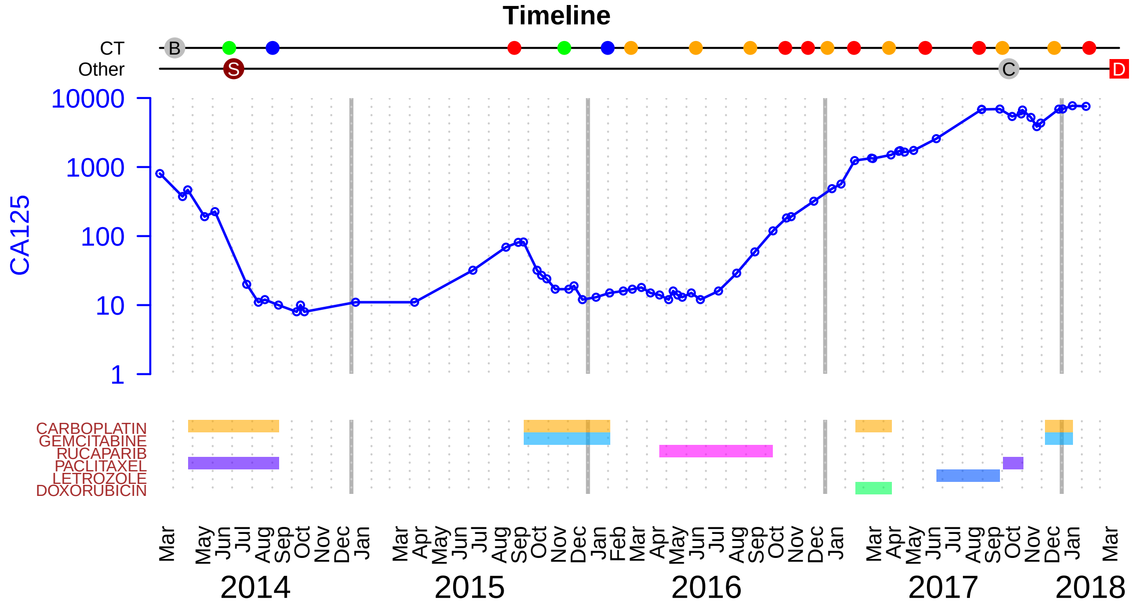


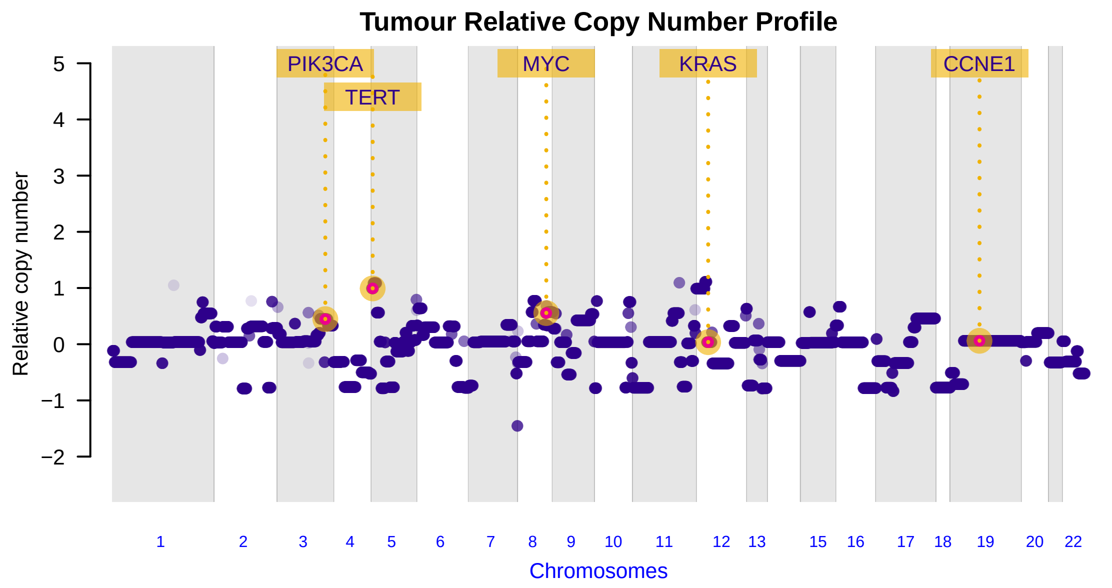


#### Spheroids drug response

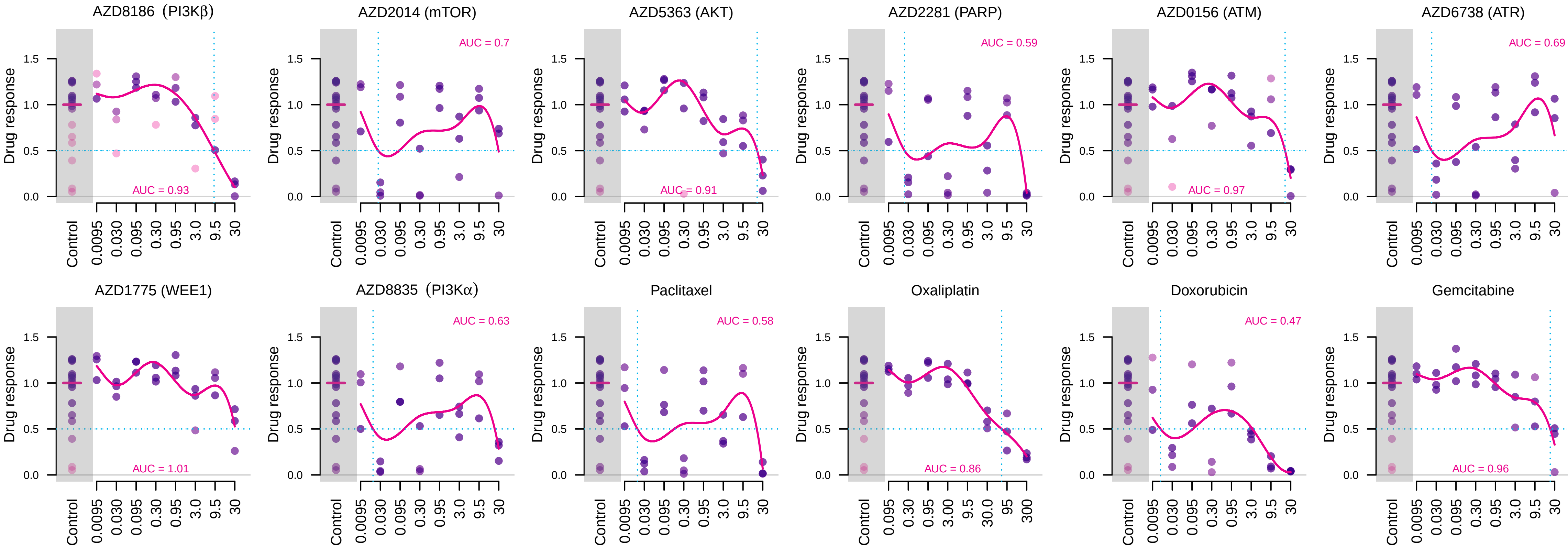

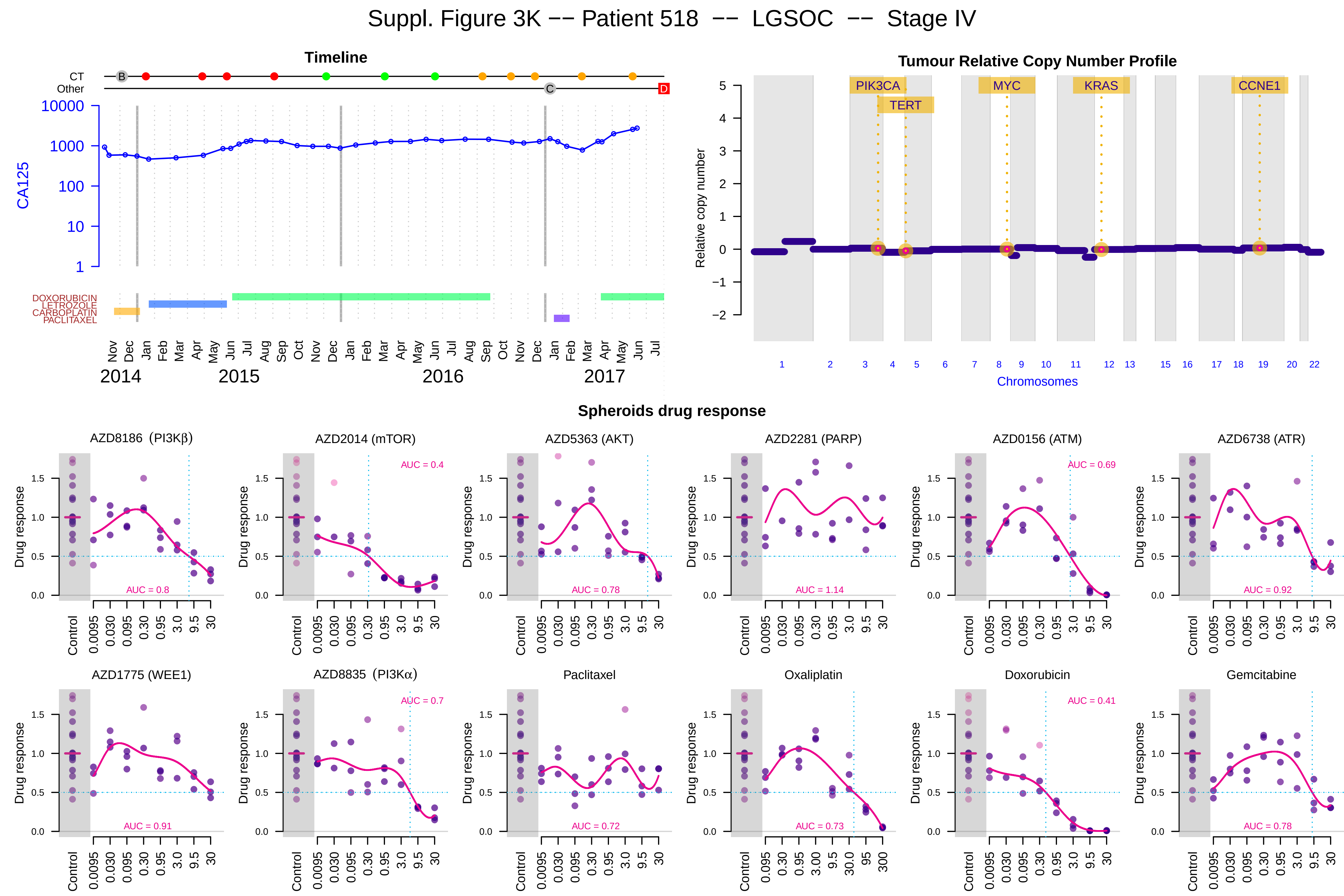
Suppl. Figure 3K -- Patient 518 -- LGSOC -- Stage IV
Timeline
CT
Other
CA125
DOXORUBICIN
LETROZOLE
CARBOPLATIN
PACLITAXEL
2014
2015
2016
2017
Tumour Relative Copy Number Profile
Relative copy number
PIK3CA
TERT
MYC
KRAS
CCNE1
Chromosomes
Spheroids drug response
AZD8186 (PI3Kβ)
AZD2014 (mTOR)
AZD5363 (AKT)
AZD2281 (PARP)
AZD0156 (ATM)
AZD6738 (ATR)
AZD1775 (WEE1)
AZD8835 (PI3Kα)
Paclitaxel
Oxaliplatin
Doxorubicin
Gemcitabine
Drug response
Control
AUC = 0.8
AUC = 0.4
AUC = 0.78
AUC = 1.14
AUC = 0.69
AUC = 0.92
AUC = 0.91
AUC = 0.7
AUC = 0.72
AUC = 0.73
AUC = 0.41
AUC = 0.78

#### Suppl. Figure 3L -- Patient 525 -- HGSOC -- Stage IV

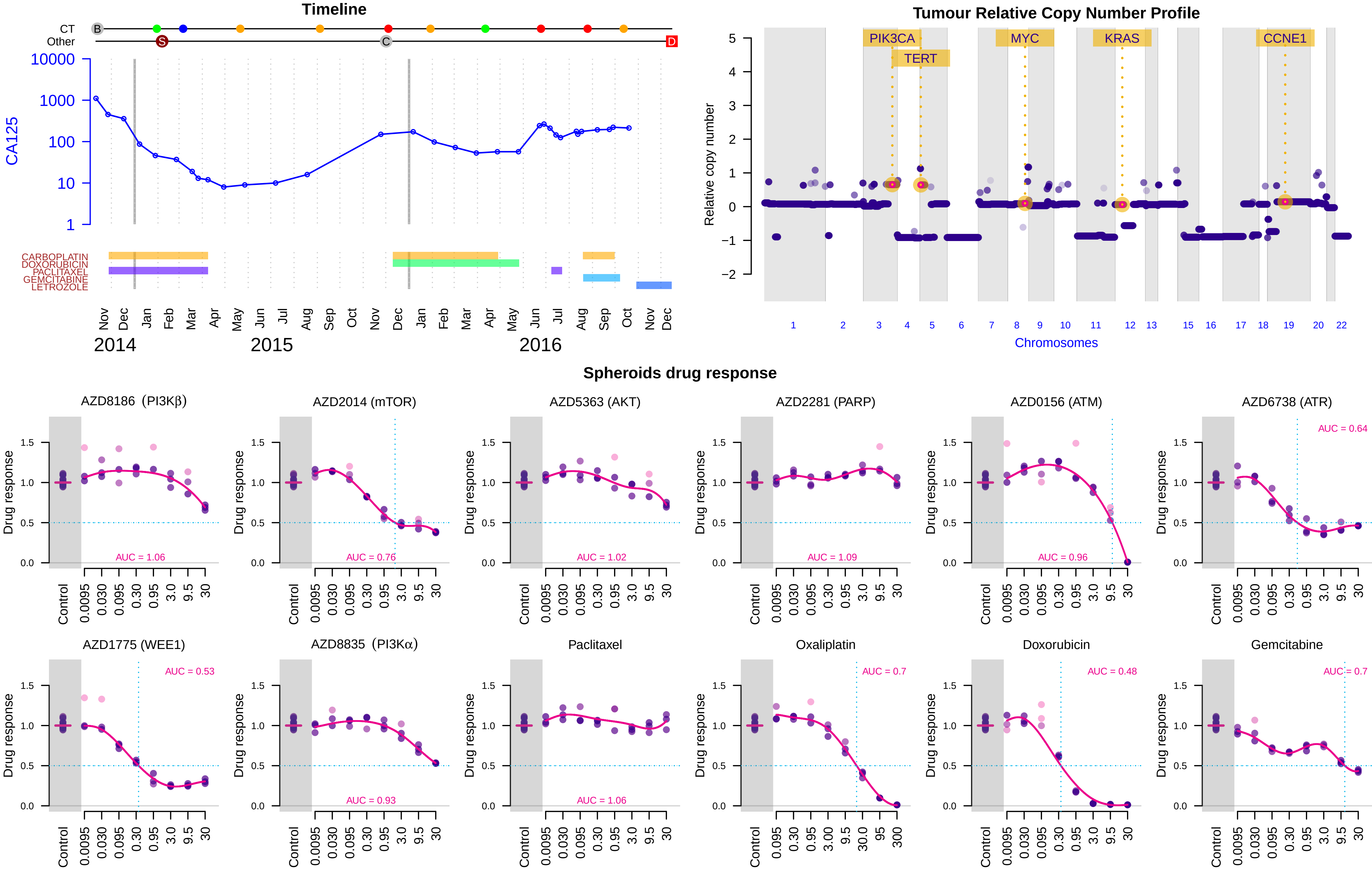

Suppl. Figure 3M -- Patient 626 -- HGSOC -- Stage III

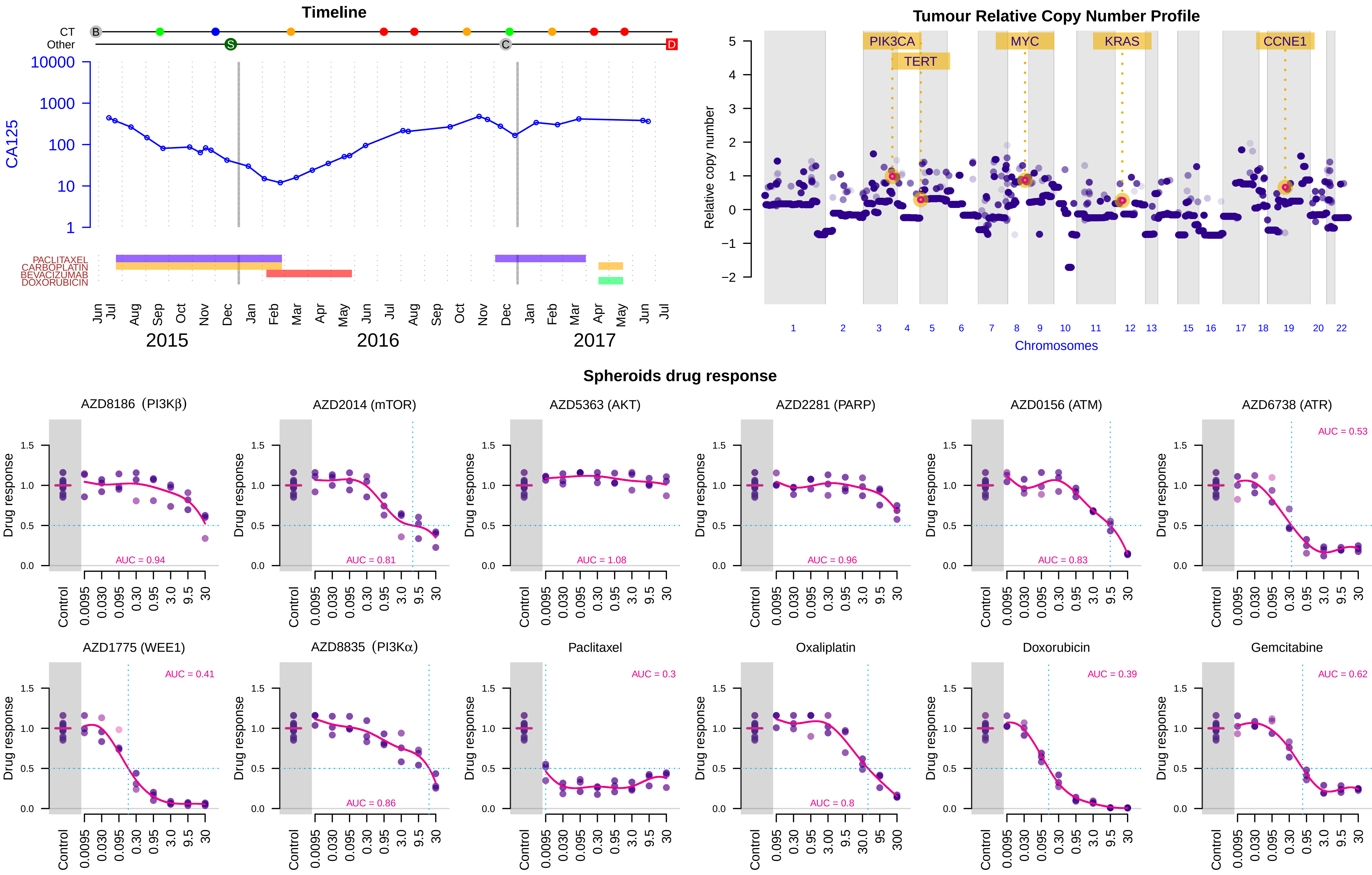

Suppl. Figure 3N -- Patient 648 -- HGSOC -- Stage III

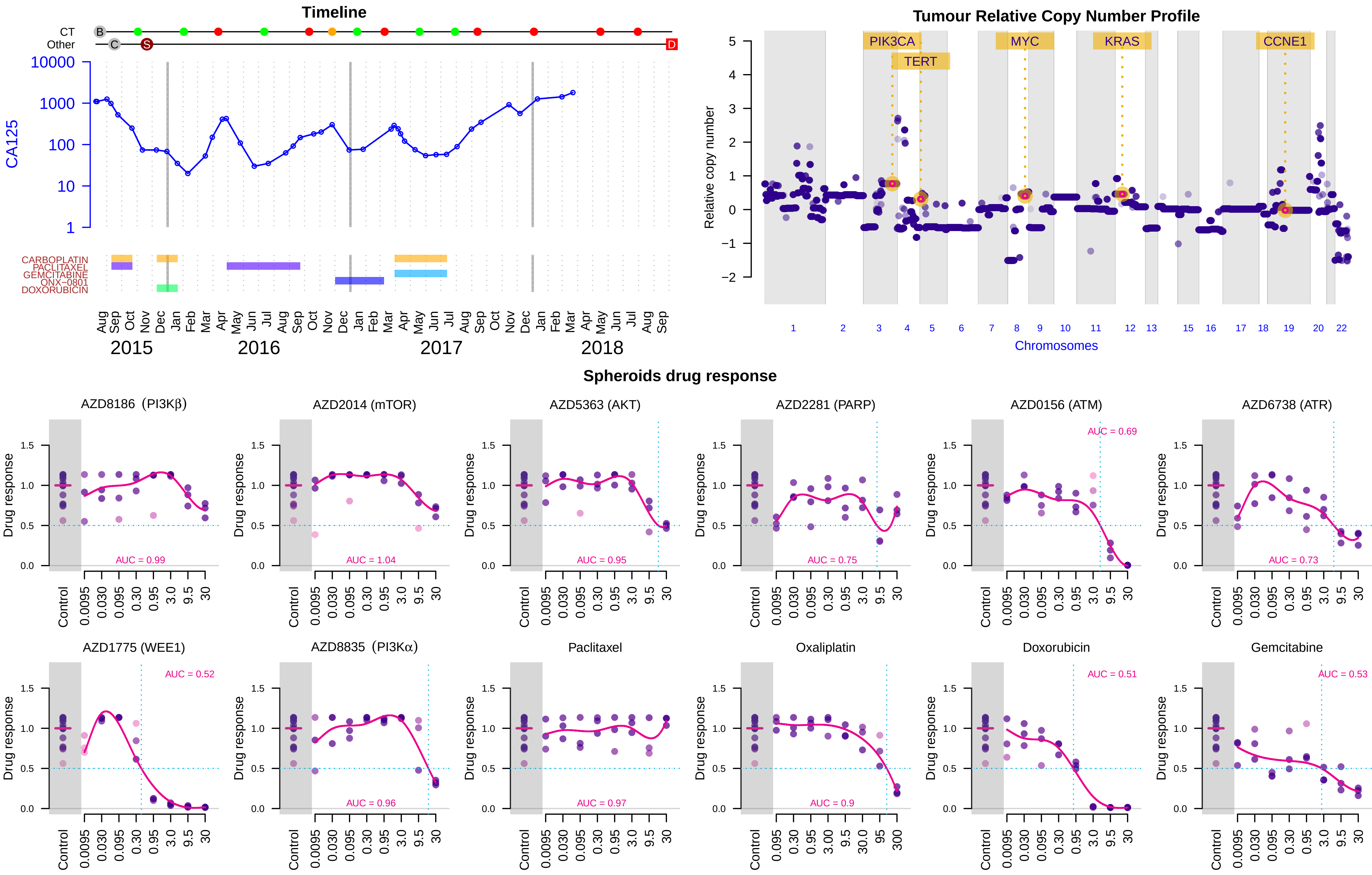

### Suppl. Figure 3O -- Patient 669 -- HGSOC -- Stage IV

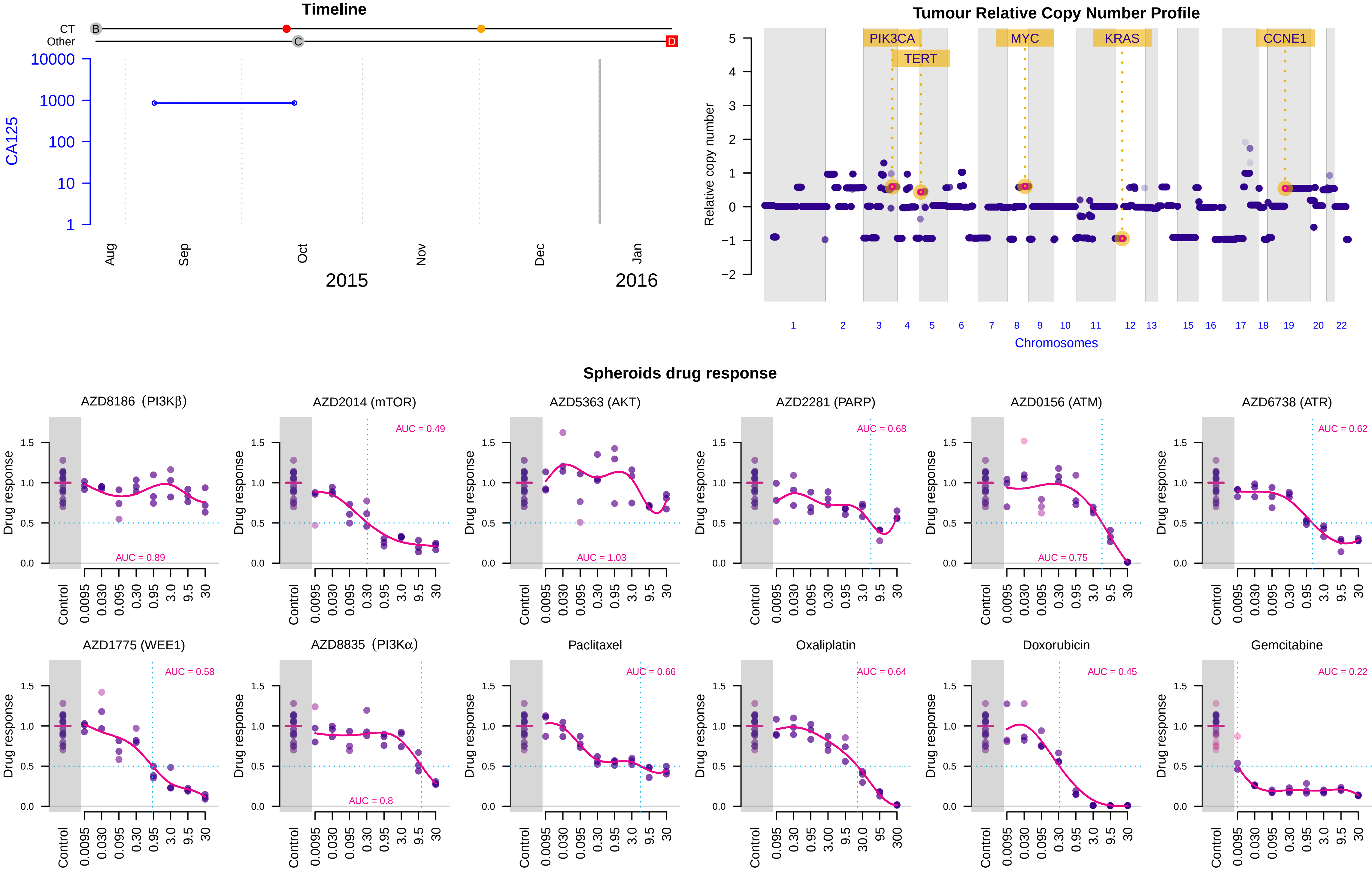

Suppl. Figure 3P -- Patient 680 -- CCOC -- Stage III

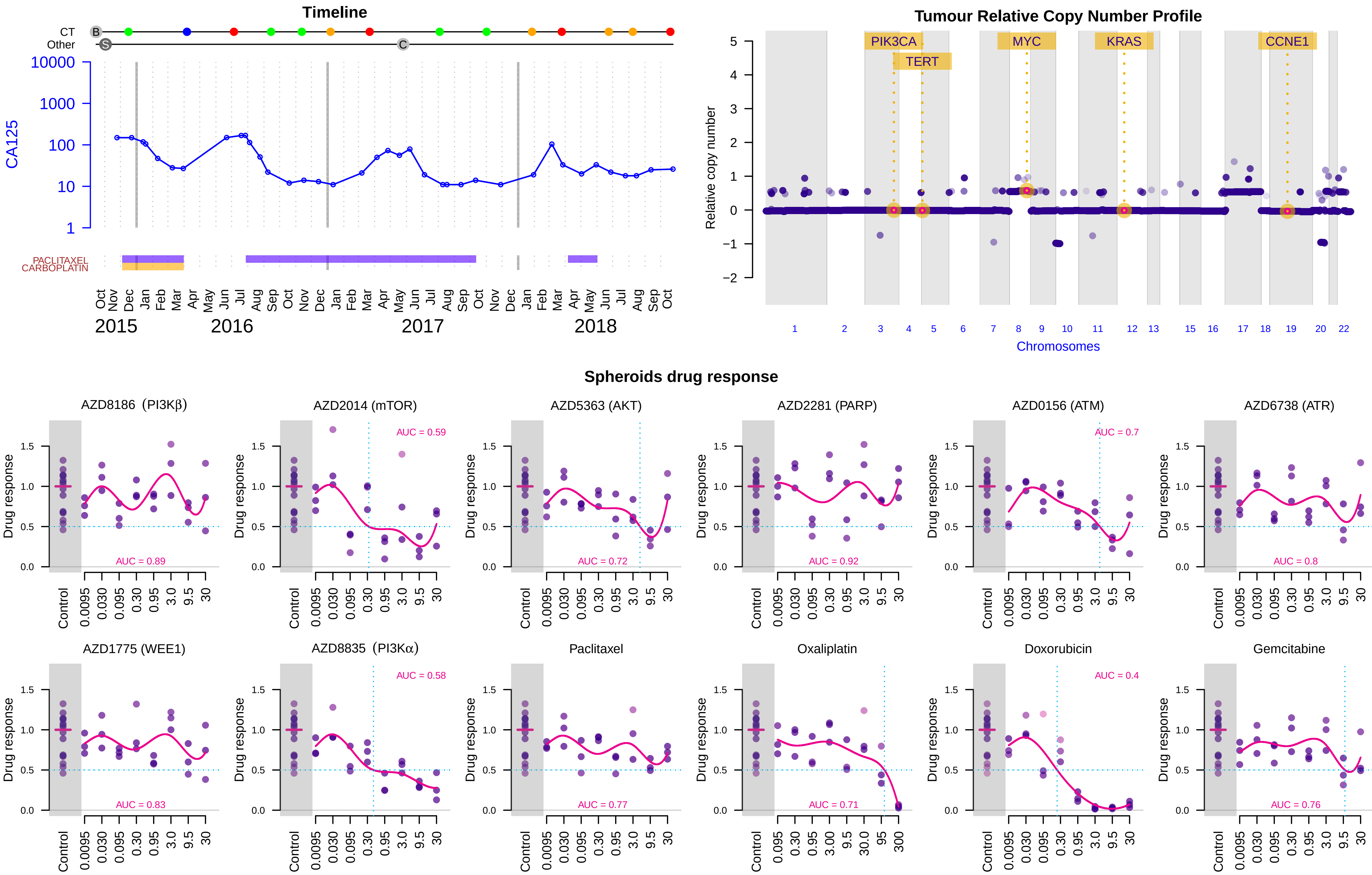

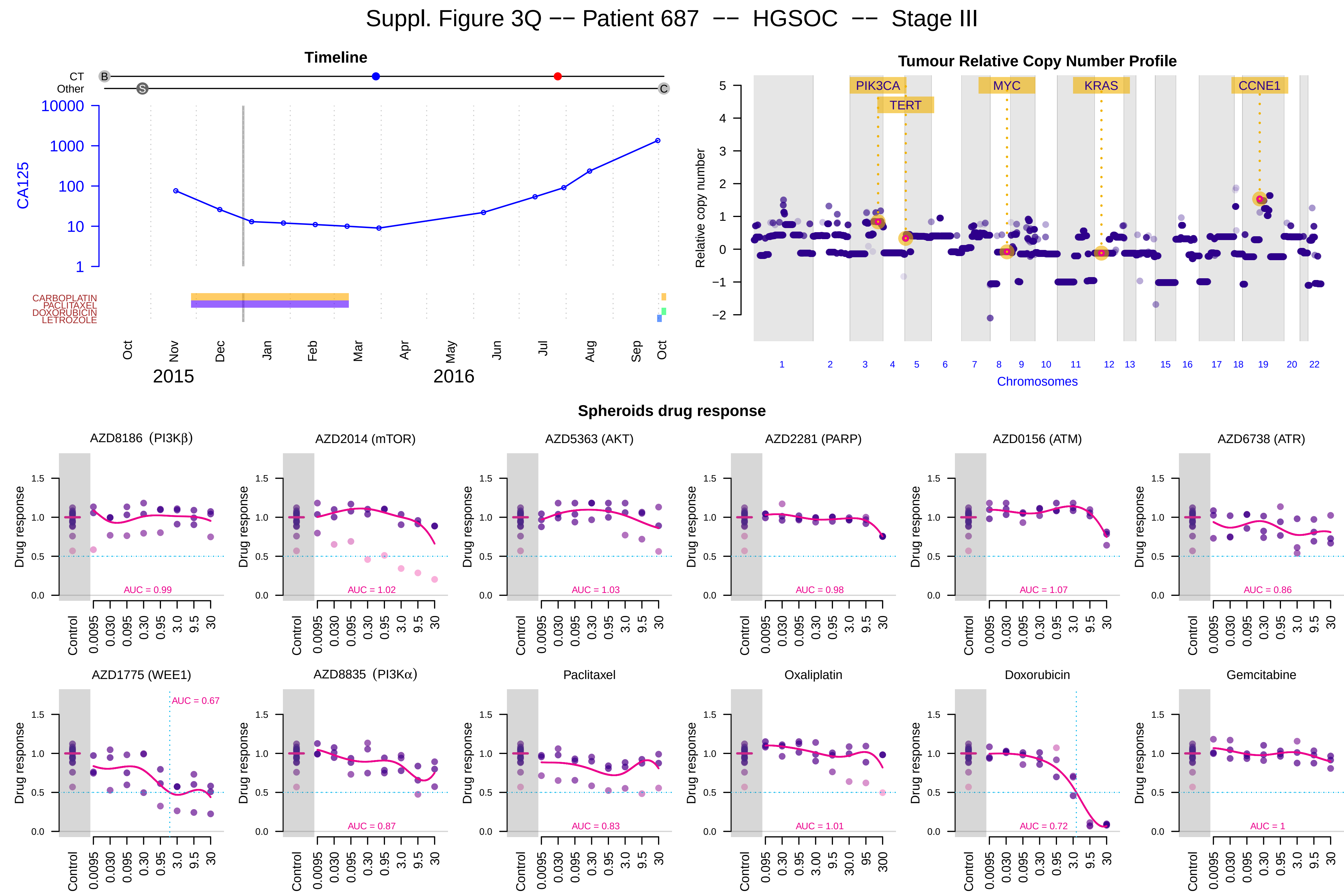

Suppl. Figure 3Q -- Patient 687 -- HGSOC -- Stage III
Timeline
CT
Other
B
S
C
CA125
10000
1000
100
10
1
CARBOPLATIN
PACLITAXEL
DOXORUBICIN
LETROZOLE
Oct
Nov
Dec
Jan
Feb
Mar
Apr
May
Jun
Jul
Aug
Sep
Oct
2015
2016
Tumour Relative Copy Number Profile
Relative copy number
PIK3CA
TERT
MYC
KRAS
CCNE1
Chromosomes
Spheroids drug response
AZD8186 (PI3Kβ)
AUC = 0.99
AZD2014 (mTOR)
AUC = 1.02
AZD5363 (AKT)
AUC = 1.03
AZD2281 (PARP)
AUC = 0.98
AZD0156 (ATM)
AUC = 1.07
AZD6738 (ATR)
AUC = 0.86
AZD1775 (WEE1)
AUC = 0.67
AZD8835 (PI3Kα)
AUC = 0.87
Paclitaxel
AUC = 0.83
Oxaliplatin
AUC = 1.01
Doxorubicin
AUC = 0.72
Gemcitabine
AUC = 1
Drug response
Control

### Suppl. Figure 3R -- Patient 720 -- HGSOC -- Stage III

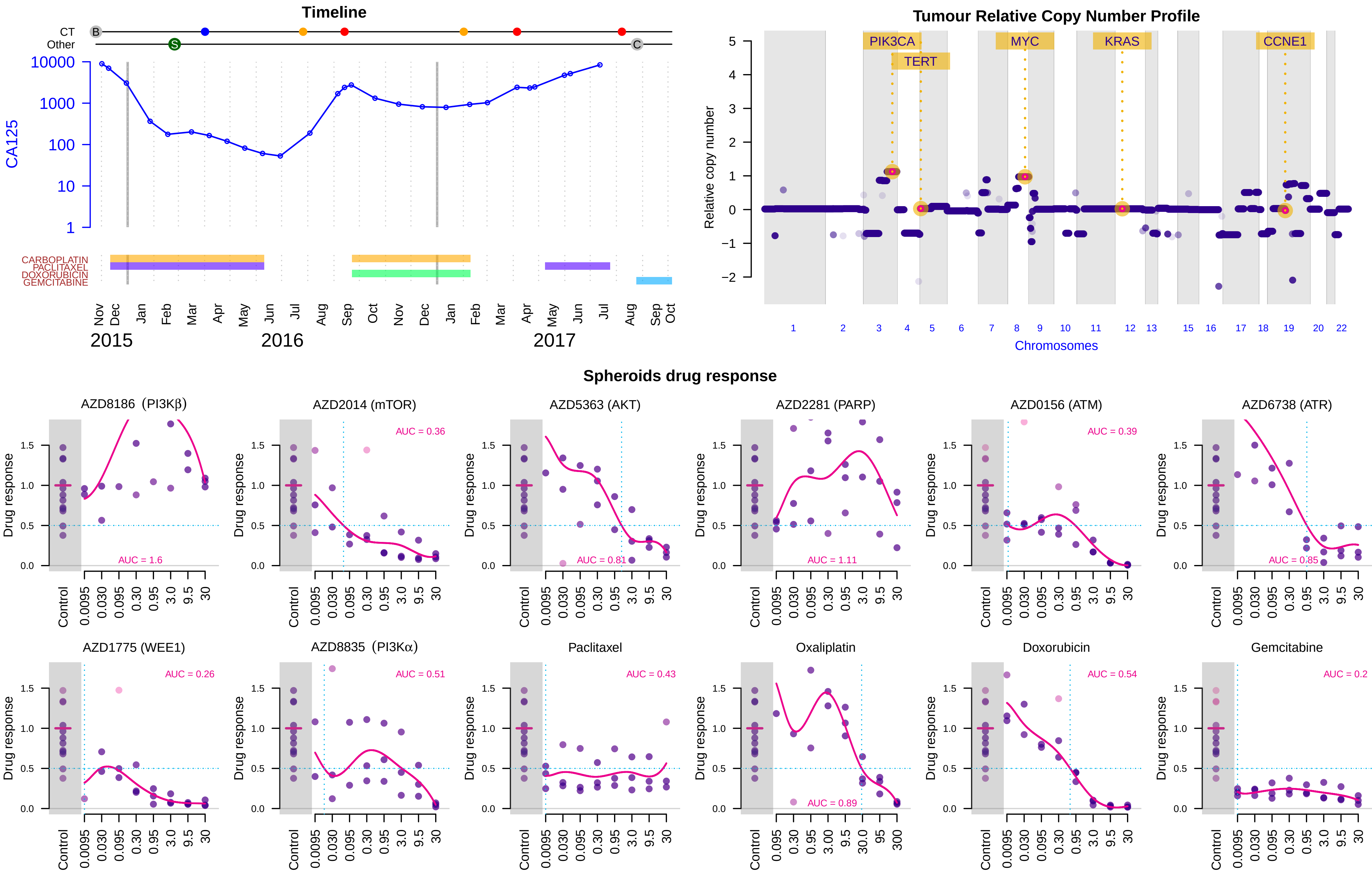

Suppl. Figure 3S -- Patient 783 -- HGS undifferentiated -- Stage NA

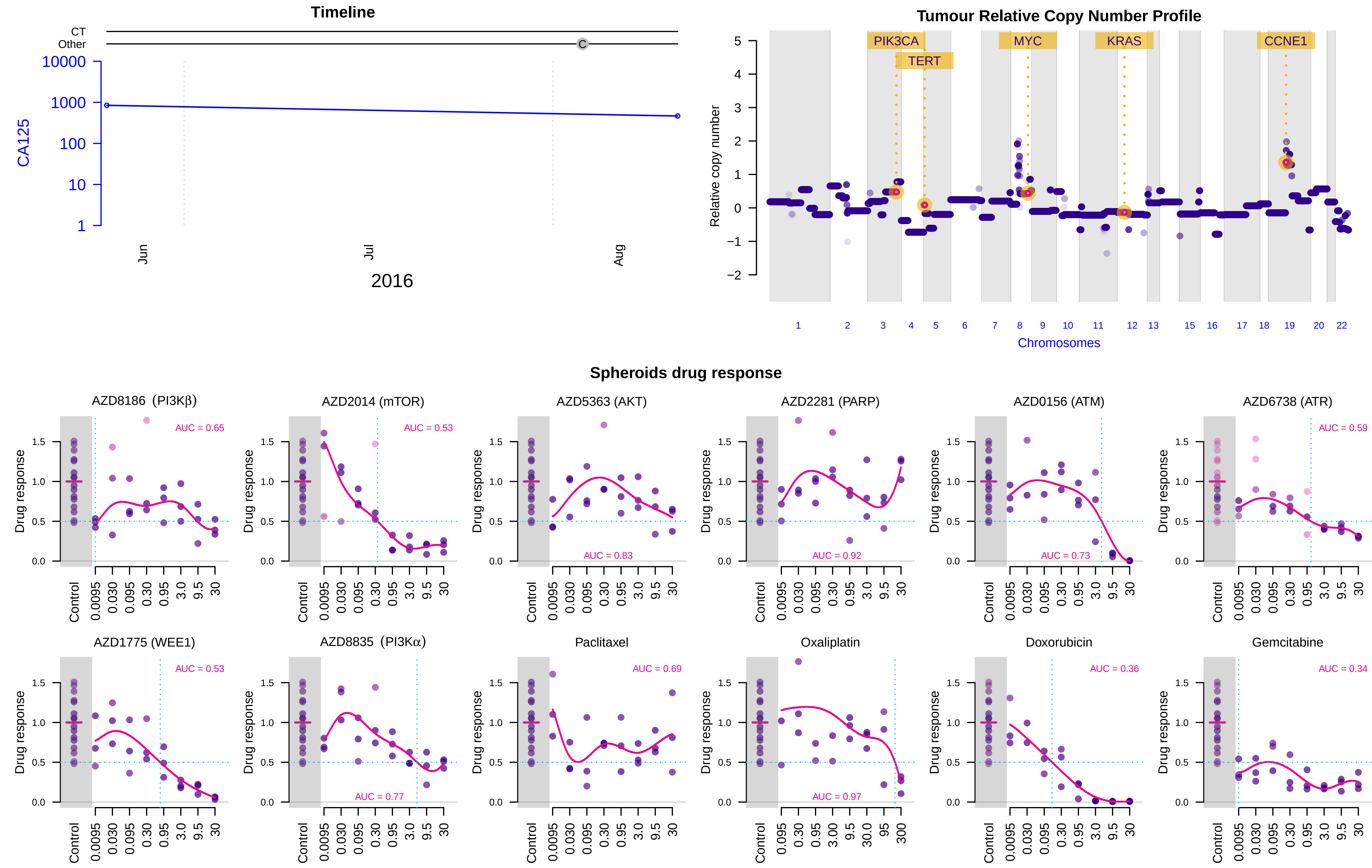

### Suppl. Figure 3T -- Patient 788 -- HGSOC -- Stage IV

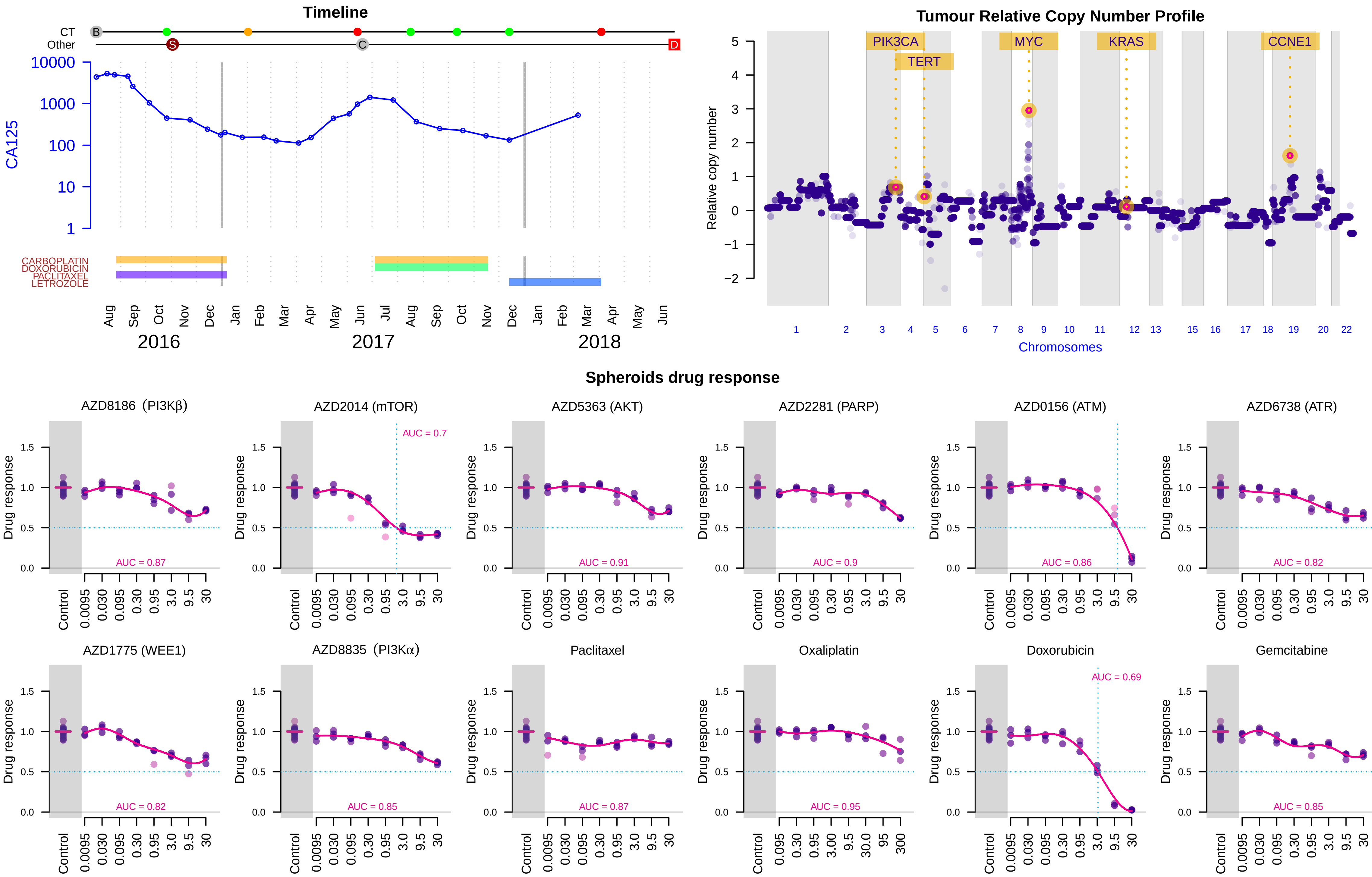

Suppl. Figure 3U -- Patient 800 -- HGS Peritoneum -- Stage III

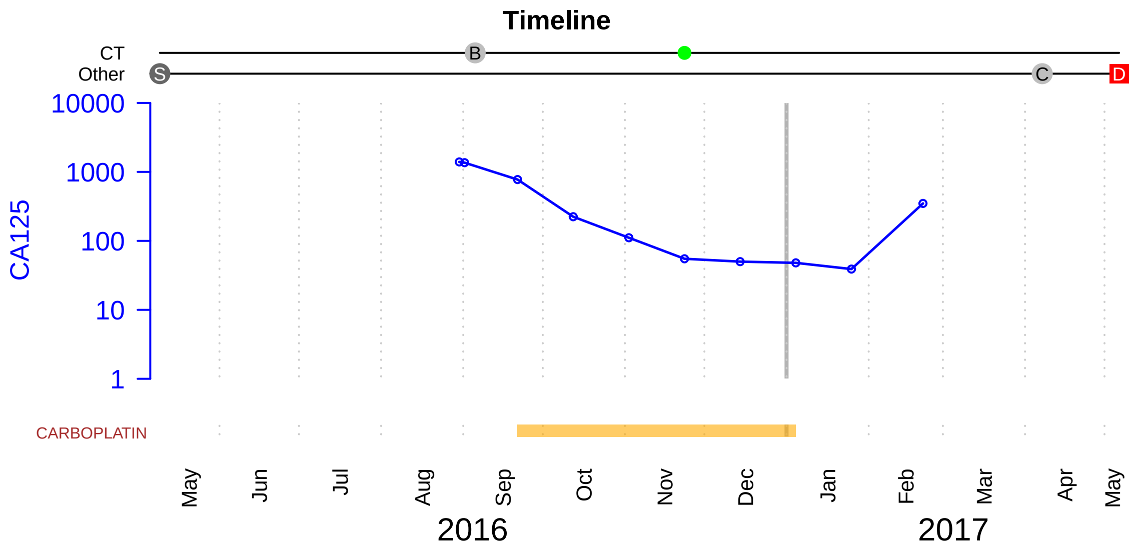


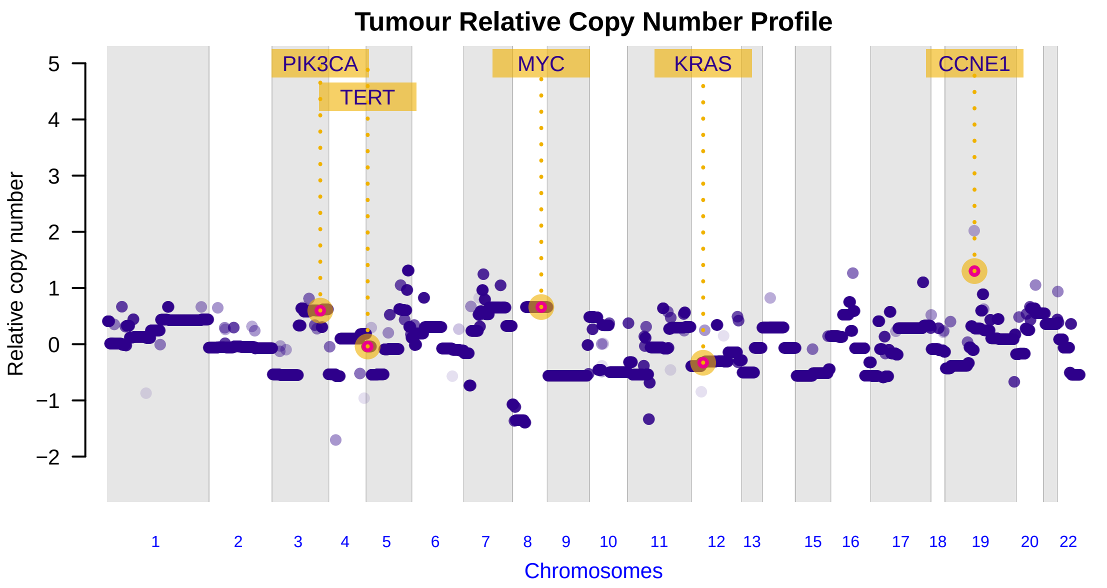


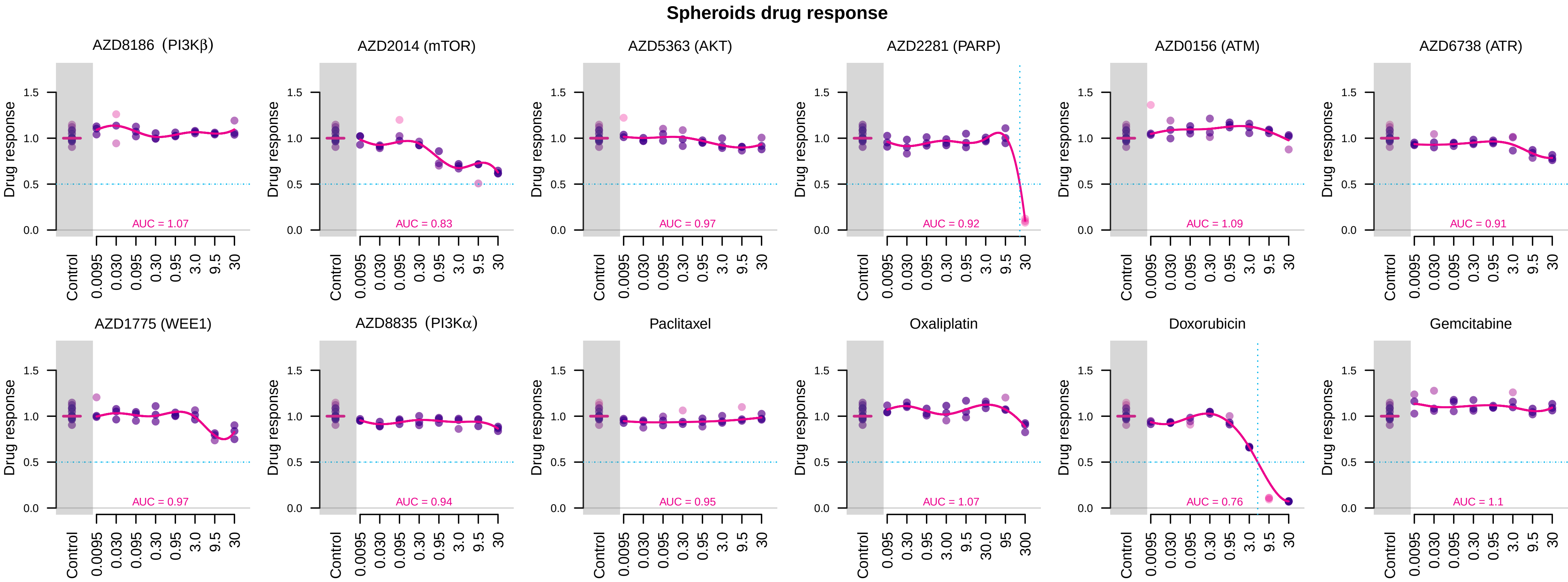

### Suppl. Figure 3V -- Patient 819 -- HGSOC -- Stage III

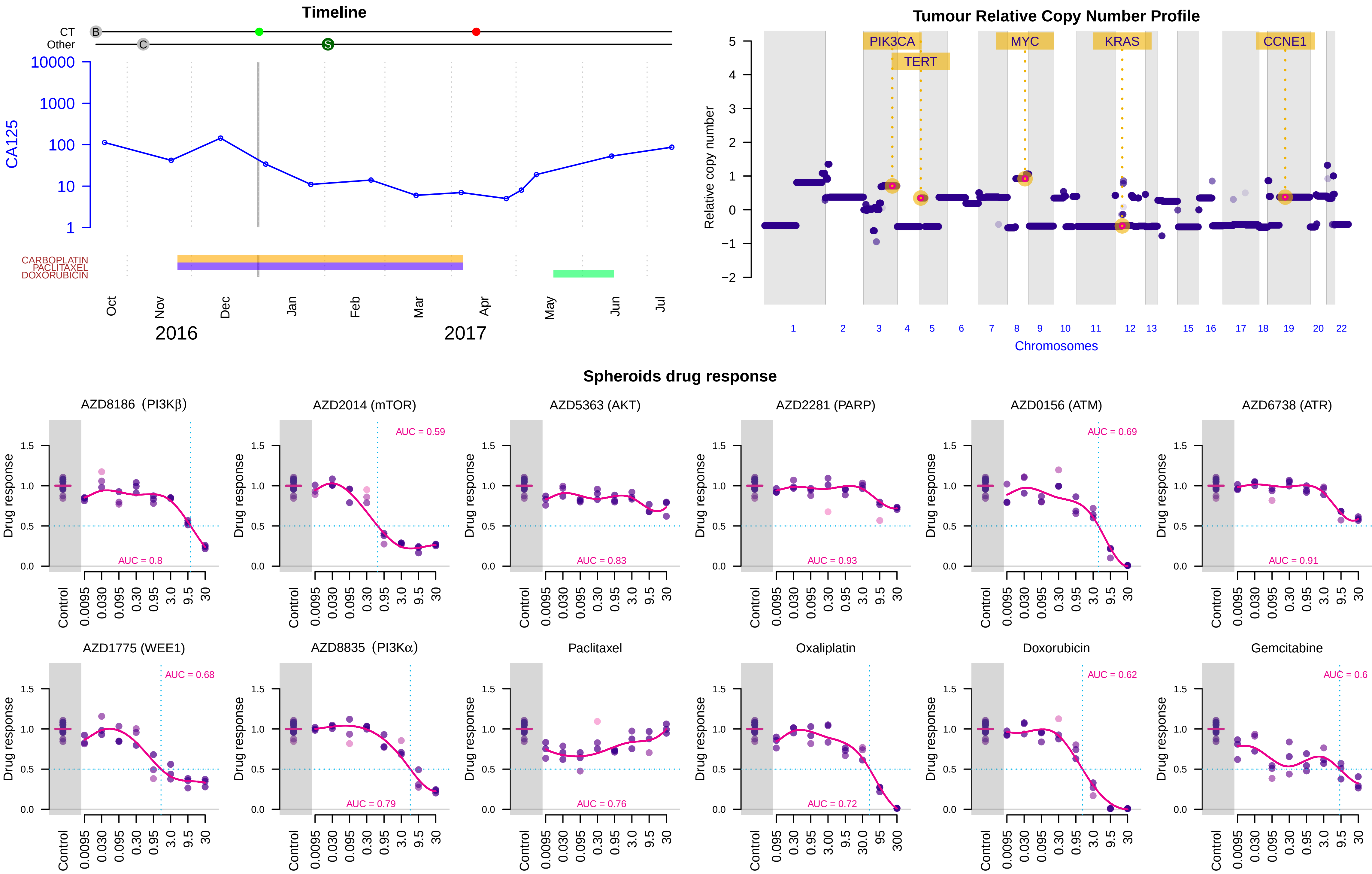

### Suppl. Figure 3W -- Patient 839 -- HGSOC -- Stage III

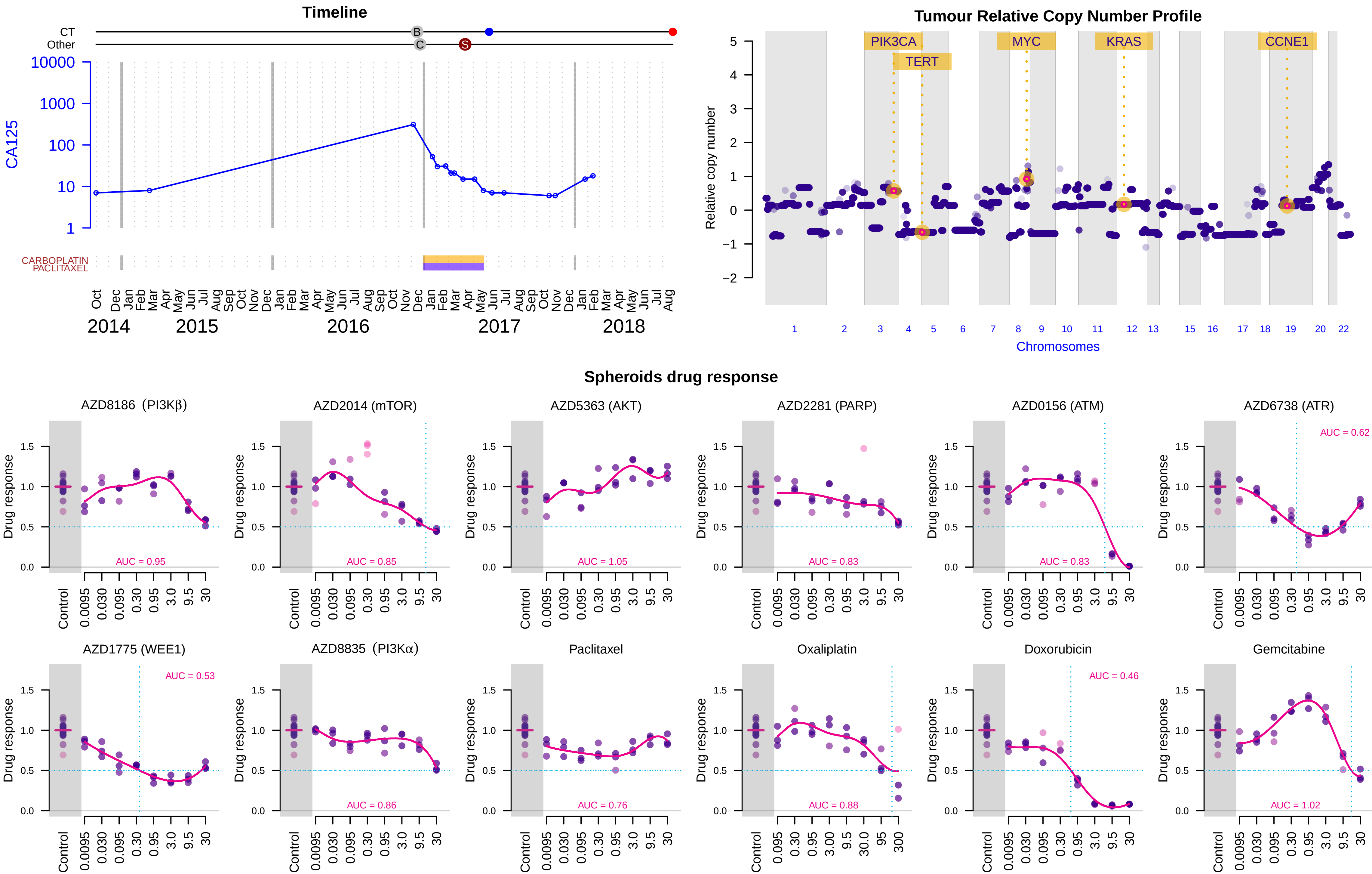

### Suppl. Figure 3X -- Patient 864 -- HGS Peritoneum -- Stage III

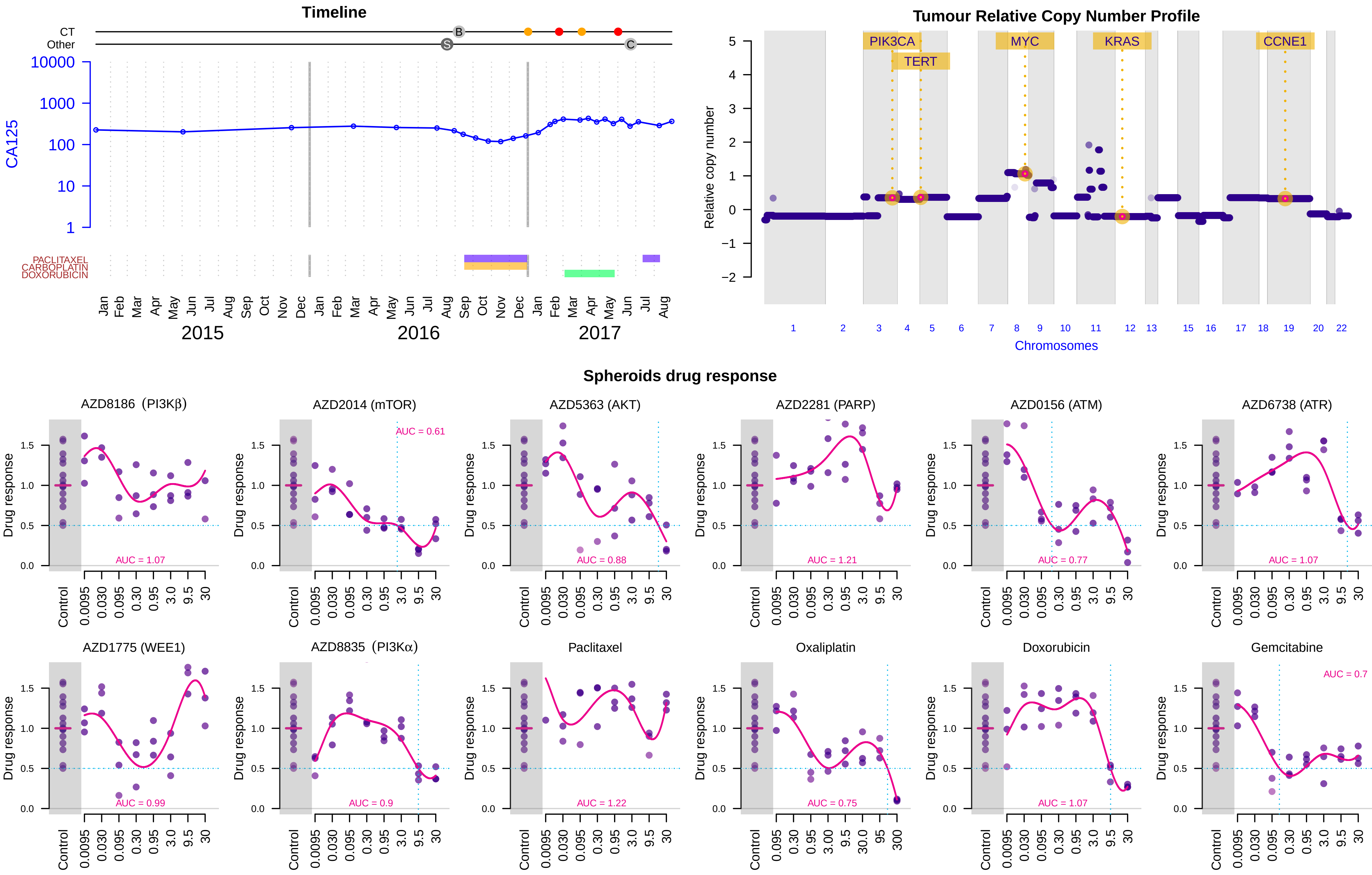

Suppl. Figure 3Y -- Patient 875 -- HGSOC -- Stage III

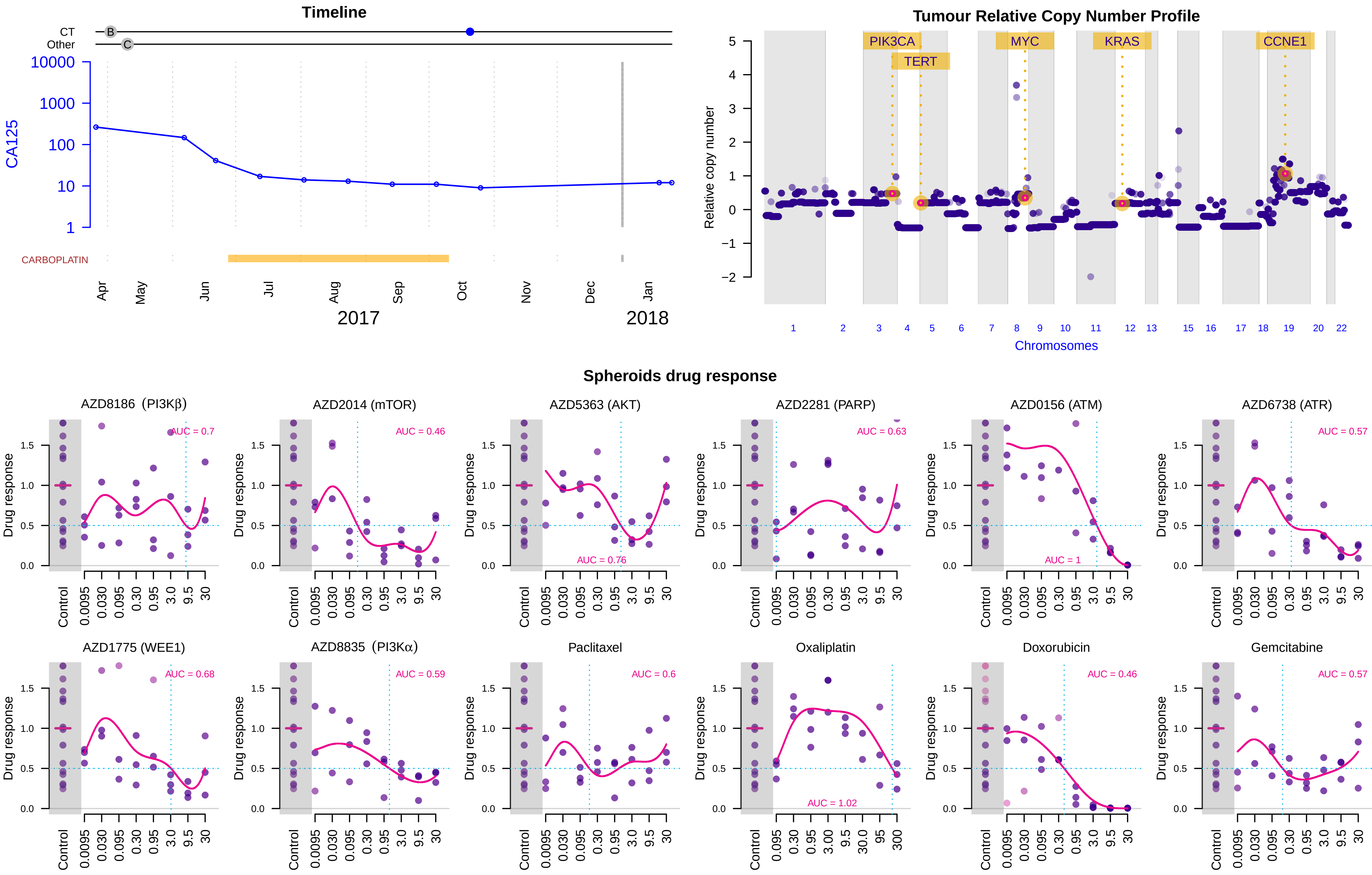

Suppl. Figure 3Z -- Patient 904 -- LGSOC -- Stage III

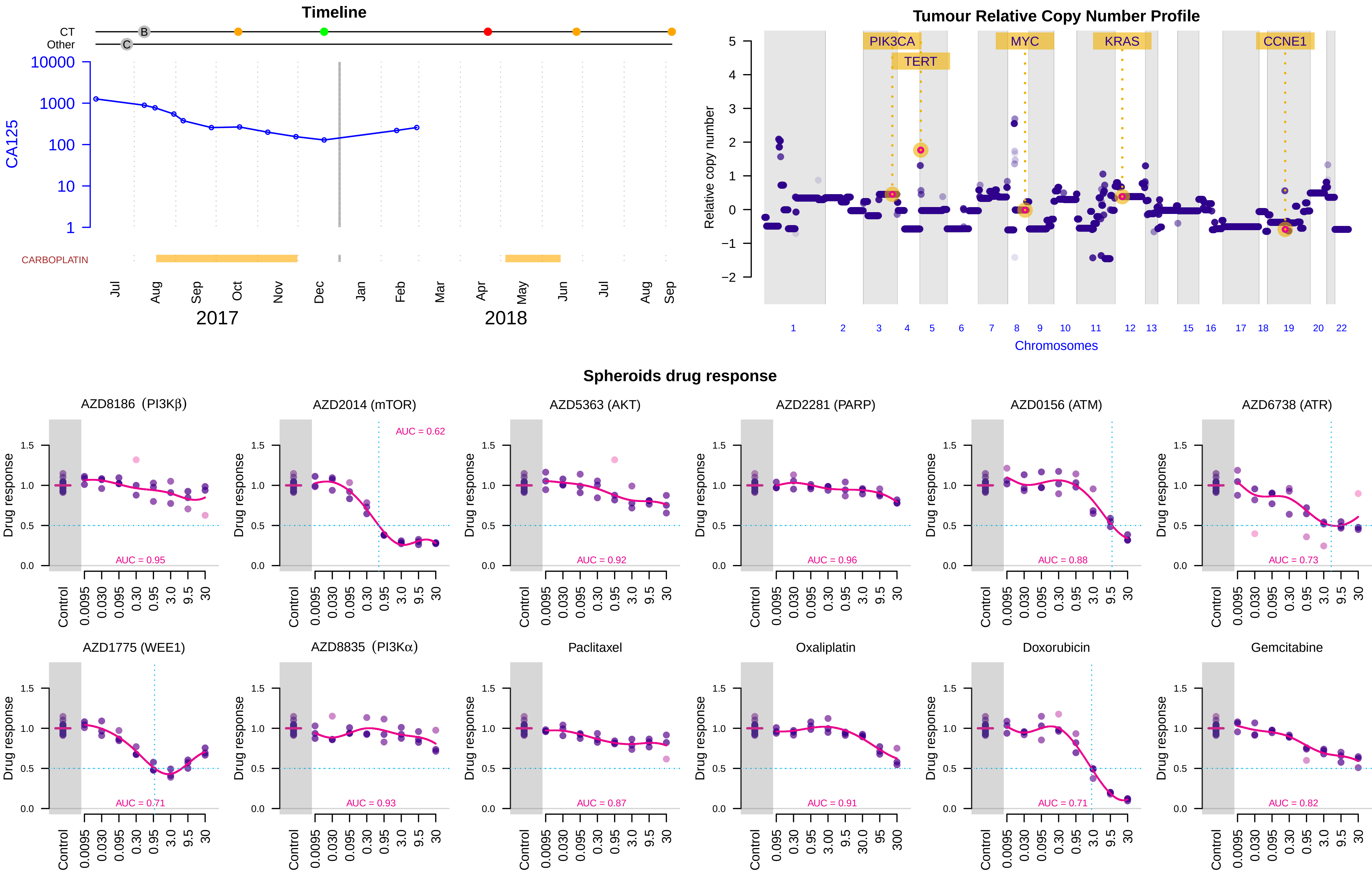
